# TMT-Opsins differentially modulate medaka brain function in a context-dependent manner

**DOI:** 10.1101/698480

**Authors:** Bruno M. Fontinha, Theresa Zekoll, Mariam Al-Rawi, Miguel Gallach, Florian Reithofer, Alison J. Barker, Maximilian Hofbauer, Ruth M. Fischer, Arndt von Haeseler, Herwig Baier, Kristin Tessmar-Raible

**Affiliations:** Max F. Perutz Laboratories, University of Vienna, Campus Vienna Biocenter, Vienna, Austria; Research Platform ‘‘Rhythms of Life,’’ University of Vienna, Vienna, Austria; Center for Integrative Bioinformatics Vienna, Max F. Perutz Laboratories, University of Vienna and Medical University of Vienna, Vienna, Austria; Bioinformatics and Computational Biology, Faculty of Computer Science, University of Vienna, Vienna, Austria; Max Planck Institute of Neurobiology, Martinsried, Germany; loopbio GmbH, Wien, Austria; FENS-Kavli network of Excellence

**Keywords:** non-visual opsins, behavior, modulation, transcriptome, teleosts

## Abstract

Vertebrate behavior is strongly influenced by light. Light receptors, encoded by functional Opsin proteins, are present inside the vertebrate brain and peripheral tissues. This expression feature is present from fishes to human and appears to be particularly prominent in diurnal vertebrates. Despite their conserved widespread occurrence, the non-visual functions of Opsins are still largely enigmatic. This is even more apparent when considering the high number of Opsins. Teleosts possess around 40 Opsin genes, present from young developmental stages to adulthood. Many of these Opsins have been shown to function as light receptors. This raises the question, if this large number might mainly reflect functional redundancy or rather maximally enables teleosts to optimally use the complex light information present under water. We focus on *tmt-opsin1b* and *tmt-opsin2*, c-Opsins with ancestral-type sequence features, conserved across several vertebrate phyla, expressed with partly similar expression in non-rod, non-cone, non-RGCs brain tissues and a similar spectral sensitivity. The characterization of the single mutants revealed age- and light-dependent behavioral changes, as well as an impact on the levels of the preprohormone *sst1b* and the voltage-gated sodium channel subunit *scn12aa.* The amount of day-time rest is affected independently of eyes, pineal and the circadian clock in *tmt-opsin1b* mutants. We further focused on day-time behavior and the molecular changes in *tmt-opsin1b/2* double mutants, and revealed that – despite their similar expression and spectral features– these Opsins interact in part non-additively. Specifically, double mutants complement molecular and (age-dependently) behavioral phenotypes observed in single mutants.Our work provides a starting point to disentangle the highly complex interactions of vertebrate non-visual Opsins, suggesting that *tmt-opsin*-expressing cells together with other visual and non-visual Opsins provide detailed light information to the organism for behavioral fine-tuning. This work also provides a stepping stone to unravel how vertebrate species with conserved Opsins, but in different ecological niches respond to similar light cues and how human generated artificial light might impact on behavioral processes in natural environments.

## Introduction

Organisms are exposed to a large range of light intensities and spectral changes. While humans are well aware of the visual inputs from their environments, the range of intensity and spectral changes that animals, including humans, naturally undergo are less consciously experienced. Across the day, light intensity routinely differs by several orders of magnitude depending on the angle of the sun above the horizon and weather conditions (e.g. a 10^3^-fold difference occurs between a sunny day and the moments before a thunderstorm, which is a similar difference between a sunny day and the average office illumination)[1].

While vision evolved multiple mechanisms to compensate for these differences, allowing us to see almost equally well across about 10^6^ orders of magnitude of light intensity [1], light differences have wider effects on physiology and behavior than solely impacting vision. This has been well documented for vertebrates, including mammals [2]. It prominently affects human (and other mammalian) physiology, mood and cognitive function via the entrainment of the circadian clock [3, 4]. In mouse it was shown that in addition to the circadian clock setting, light information conveyed via the perihabenular nucleus also directly affects brain regions that control mood [5], while moderate exposure to UVB light promotes the biosynthesis of glutamate and results in enhanced learning and memory [6]. Furthermore, photoperiodic effects on mammalian mood, cognition and fear behavior are well documented, which are for example connected with changes in transcript levels of *tyrosine hydroxylase* and *pre-prosomatostatin1* [7].

These far-reaching impacts of light on behavior and physiology are not just documented for mammals, but spanning across vertebrates. Seasonality, and in particular photoperiod, controls major hormonal changes connected to breeding, molt, and song production in birds (reviewed in [8]), while light modulates motor behavior independently of the eyes in frogs [9] and teleosts [10].

The past years revealed that Opsins, the proteins mediating light sensation in vertebrates, are present in various cell types inside and outside the vertebrate eyes, including specific brain neurons [11-14]. Zebrafish possess 42 *opsin* genes [15], and additional genomic and transcriptomic data suggest that *opsin* gene numbers are similarly high in other teleost species (http://www.ensembl.org).

Biochemical analyses, tissue culture assays and electrophysiological recordings on brain tissue suggest that most, if not all, of these Opsins can function as light receptors [14-18]. Light has been shown to reach deep brain regions of several mammals and birds [11, 19, 2]. Given the typically smaller sizes of fish brains, especially medaka and zebrafish, it is highly conceivable that light will reach cells inside the fish brain. Based on the expression of several Opsins in inter- and motorneurons in larval, as well as the adult zebra- and medaka fish brains we suggested that these Opsins could modulate information processing, depending on ambient light conditions [14].

Studies on young zebrafish larvae indeed implicate non-visual Opsins in specific light-dependent behaviors, such as the photomotor response present at 30 hours-post-fertilization (hpf) [20], for which however the exact identity of the Opsin(s) involved remained unclear. The suppression of spontaneous coiling behavior in larvae younger than 24 hpf is for its green light component mediated by VAL-opsinA [21], while melanopsins have indirectly been implicated to mediate part of the dark photokinesis response (in 5-7 days-post-fertilization (dpf) old larvae) [13]. The latter assay was further extended to investigate search pattern strategies of 6-7 dpf old larvae under extended periods of sudden darkness. In this assay Cas9/Crispr engineered *opn4a* mutants showed altered local search behavior [22]. Combinational analyses of enucleated wildtypes (wt) fish*, otpa* mutants and enucleated *otpa* mutants indicate a contribution of brain-expressed *opn4a* [22]. However, as there was no comparison between enucleated *opn4a* mutant and wt fish, the contribution of *opn4a* expressed in the eye versus the brain [23] remained unresolved.

While these studies have started to provide insights into the understanding of non-visual/ deep brain Opsin functions, several important biological questions have remained entirely unaddressed.

First, non-visual Opsins are not just present during young larval stages, and their later functions remain unclear. How do the early functions compare to functions during later, juvenile stages, when the nervous system has further grown and differentiated?

Second, another line of evidence suggests that non-visual Opsins are involved in conveying light information for chronobiological functions, such as the circadian clocks autonomously present in different fish tissues [24, 25]. Outside the tropical/ sub-tropical zebrafish, non-visual Opsins have also been suggested to convey photoperiod information, particularly to the reproductive system [26]. Is there evidence for such a function in a species for which seasonal cues are naturally relevant?

Third, one of the maybe most puzzling question concerns the number of Opsins in teleost fish. Why are there so many? Is it simply redundancy of the system or do these opsins control physiology and/or behavior in a more complex, possibly synergistic manner?

Here we focus on the functional characterization of two members of the Encephalopsin/TMT-Opsin (ETO) family, which is characterized by a particularly low rate of sequence changes over time and represents ancestral c-Opsins [14, 27]. One of its subfamilies (Encephalopsin) is conserved up to placental mammals [14, 28, 29]. We focused our analyses on medakafish, since these fish show a light-dependent seasonal breeding response as an adaptation to the photoperiod changes naturally occurring in their habitat [30, 31]. We decided to specifically focus on Tmt-opsin1b and Tmt-opsin2. Spectral analyses in cells and on purified proteins suggest that these Opsins have a similar wavelength preference, while also being expressed in close vicinity or even partly overlappingly ([14, 32] and this study), thus providing an interesting entry point for testing complementing vs. non-complementing functions. We generated medaka *tmt-opsin1b* and *tmt-opsin2* mutants, and analyzed mutant and sibling wildtype fish in multiple behavioral and molecular assays. Our functional work on even just two non-visual Opsins already indicates a complex, (at least in part) non-redundant light information processing system, which modulates a neurohormone and a voltage-gated sodium-channel subunit, as well as behavior. In addition, our work provides several examples how relatively small differences (in age and light intensity) manifests themselves in significant behavioral changes.

## Results

### *Ola-tmt-opsin1b* mutants exhibit light-dependent altered avoidance responses

We started our investigation with medaka *tmt-opsin1b*, by generating several independent mutant alleles using TALEN technology (Fig 1A). All three mutant alleles are predicted to result in an N-terminally truncated protein, prior to the first transmembrane domain (S1A-D,G Fig) and are thus likely functional null mutants. We confirmed the absence of functional protein with a previously generated antibody against *Ola*-TMTopsin1b [14] (Fig 1B). We next tested for noticeable developmental alterations by detailed staging of the developing larvae according to morphological criteria [33]. We did not observe any difference between wt and homozygous mutant larvae (S2A-C Fig). We also crossed a strain expressing GFP under the control of an ath5 enhancer fragment in the developing retinal ganglion cells [34, 35] with the *tmt-opsin1b*−/− fish. The resulting heterozygous, GFP+ carriers were incrossed and their larval offspring analyzed for any deviations in the on-set of GPF expression (indicative of the timing of RGC differentiation), axonal outgrowth and connectivity to the tectum (indicative of RGC and tectal development). After hatching (7-9dpf) the fish were genotyped. As we again did not observe any differences between mutant and wildtype fish (n: wt=19 ; heterozygous mutant = 27, homozygous mutant = 18, S2D-G Fig), we concluded that especially also eye and tectal development is unaffected by the *tmt-opsin1b* mutation.

**Fig 1:**
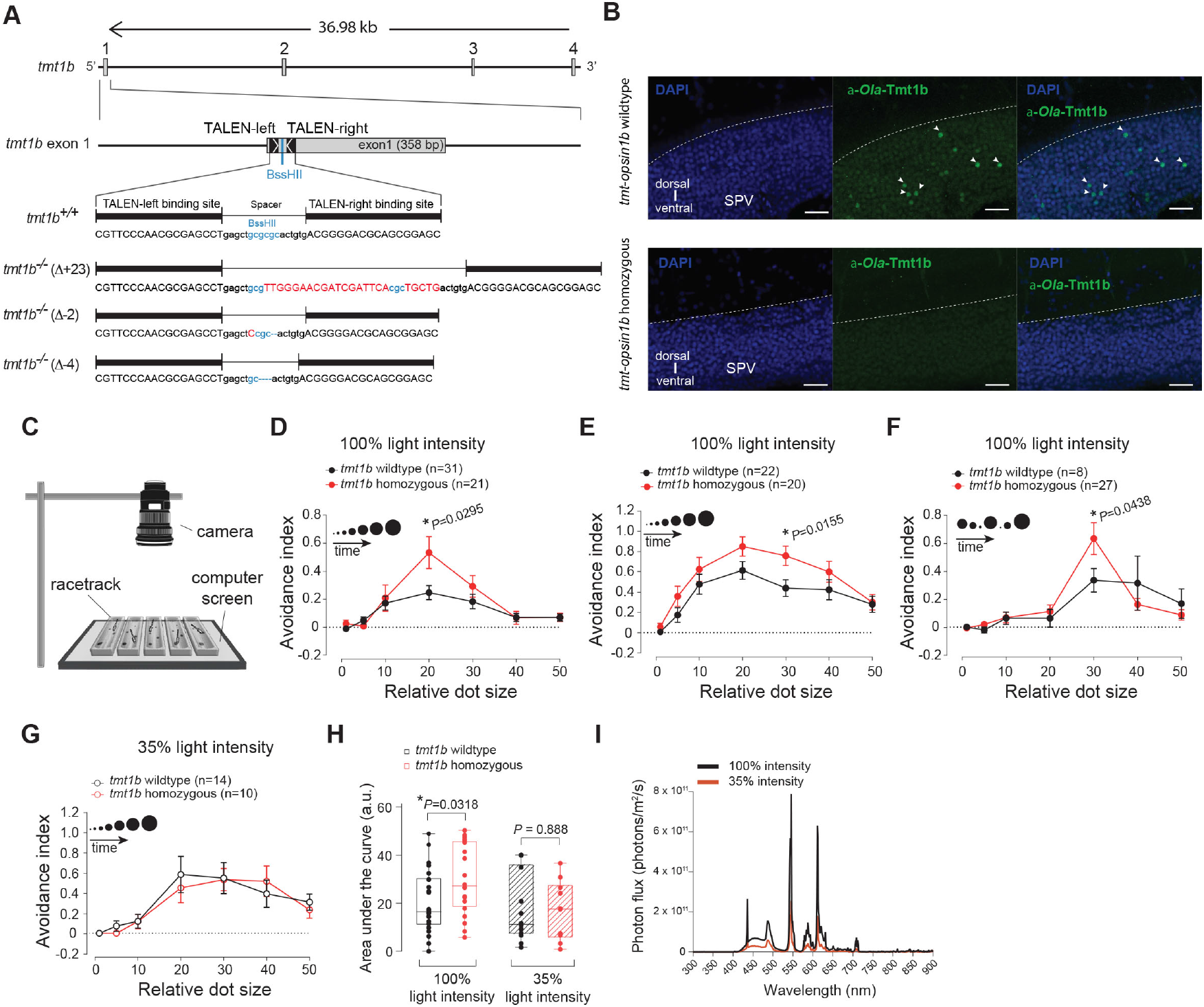
Mutations in *Ola-tmt-opsin1b* cause light-dependent differences in an avoidance response test. (**A**) Genomic locus of the Ola-tmt-opsin1b gene and corresponding mutant alterations. TALEN binding sites: black boxes; Exons: grey boxes; Inserted/substituted nucleotides: red; deleted nucleotides: “−”. (**B**) Confocal images of anti-Ola-TMT-1b (green) and nuclear DAPI (blue) staining in wildtype and tmt-opsin1b (D-2/D-2) mutant coronal tectal slices. SPV: stratum periventriculare. Scale bars: 20 μm. (**C**) Schematic of the behavioral setup. (**D**) Size discrimination tuning curve of larvae reacting to moving dots presented in ascending size order. (**E**) Behavioral analysis as in (**D**), performed in a different laboratory with a different computer screen and surrounding (Light intensity = 2.2 × 1013 photons/m2/sec). (**F**) Avoidance responses to the dots displayed in a shuffled size manner. (**G**) Avoidance responses in the ascending dot size paradigm with background light reduced to 35% of its initial intensity (35% light intensity = 0.77 × 1013 photons/m2/sec). (**H**) Total avoidance (area under the curve) under different light intensities. (**I**) Spectra of the emitted computer screen light used for the 100% (black) and the 35% light intensity (orange). Data presentation: (D-G) mean (± s.e.m.); (H) single bar: median, box: first and third quartile, whiskers: min and max; all assays done with 9-12 dpf larvae. a.u.: arbitrary units. Metadata: S1 Data. For further details on mutants, responses to contrast changes, spectra and mutant vs. wildtype development: S1 Fig, S2 Fig, S3 Fig.

Given that *Ola*-TMTopsin1b is expressed in the medaka optic tectum and reticular formation [14] (Fig 1B, Fig 4B,D), we started our functional analysis with an assay that requires these structures. It has been shown that the responses of frogs and fish to approaching stimuli of different sizes specifically require the tectum and reticular formation [36-40]. We adapted an assay in which the response of individual animals to displayed moving dots of different sizes (mimicking a range of potential prey and predator stimuli) is recorded and analyzed (here referred to as “avoidance assay”, Fig 1C, S1 Video, S2 Video). Such type of avoidance assays have been used for toads[36], larval *Xenopus*[37], mice[41], goldfish[42] and larval zebrafish[40]. Medaka larvae of different stages were collected in the morning (ZT2-5) and subjected to the avoidance assay. Newly hatched medaka larvae are immediately free-swimming and might best be compared to zebrafish larvae at about 7-8 dpf (constantly raised at 28°C), while later obvious developmental steps proceed similarly to zebrafish. Based on the responses of the larvae to the moving dots, we calculated an avoidance index (AI) for each dot-size: AI= (total number of avoidances) - (total number of approaches) divided by (total number of dots presented). Each dot size was presented six times to each fish and care was taken to use different mutant alleles in the same trial (S1H Fig).

After blind response scoring and subsequent genotyping, we observed that *tmt-opsin1b* mutant fish exhibited a significantly elevated avoidance response compared to their wildtype siblings, particularly to dot sizes 20 degrees (*P* = 0.0295, Unpaired t test with Welch’s correction; t = 2.3; df = 26.4) and 30 degrees (*P* = 0.0155, Unpaired t-test; t = 2.5; df = 40), decreasing gradually towards bigger dots (Fig 1D,E, S1 Data). Shuffling dot size order resulted in an overall shift of the response curve, presumably due to habituation of the fish to the larger dots that appeared earlier (compare Fig 1D,F, S1 Data). Importantly, the elevated response of *tmt-opsin1b* mutants vs. wildtype siblings was maintained.

Given that different lab environments resulted in slightly different response curves of the mutant fish (Fig 1D,E, S1 Data) and that TMT-Opsin1b should function as a light receptor in wildtype, we next tested if the observed elevated response in the avoidance assay would be altered by changes of light conditions. The difference of the avoidance responses present between wildtype and *tmt-opsin1b* mutants at 100% light intensity vanishes when the light intensity is reduced (Fig 1G,H, *P* = 0.0318, Unpaired t-test; t = 2.2; df = 40, without changing the spectrum, Fig 1I, S1 Data). Specifically, when we lowered the light intensity of the white background projected by the computer screen to 35% of the original light intensity, the mutant AI-curve and the wildtype AI-curve (compare Fig 1E,G,H, S1 Data), became statistically indistinguishable (*P* = 0.888, Unpaired t-test; t = 0.14; df = 22). We also tested, if the observed changes in the behavioral responses observed between the different light intensity conditions might be due to contrast differences. For this, we analyzed the avoidance index of Cab wildtypes under different contrast conditions (S1I Fig, S7 Data) by testing the fish for their responses to dots of increasing luminance (keeping dot size constant at 20 degrees). 35% lower light levels at the computer screen correspond to a Michelson contrast of 0.93, a contrast that does not cause any observable behavioral differences compared to full light levels (S1I Fig, S7 Data), making a change of contrast an unlikely explanation for the observed drop in the avoidance index of *tmt-opsin1b* mutant fish at 35% ambient light intensity. These results suggest that *tmt-opsin1b* normally mediates constant responses, even under changing light conditions (for comparisons of the light spectra to natural light levels see S3 Fig).

### *Ola-tmt-opsin1b* mutants exhibit strong age-dependent behavioral differences to wt in response to sudden light/dark changes

The avoidance assay requires the tectum for proper responses, but it also requires normal functioning eyes. Like probably all Opsins, *tmt-opsin1b* is expressed in the eye, specifically in the amacrine layer [14]. We therefore next decided to use assays that do not necessarily require eyes. Animals typically respond to changes in ambient illumination with rapid changes in movement[43]. Changes in motor behavior in response to light in blinded and pinealectomized minnows[10], eels [44] and lamprey tails [45] provided the first evidence for extraretinal, extrapineal “photomotor” behavior in vertebrates. Furthermore, in zebrafish larvae, short periods of sudden darkness result in an increased overall activity, which has been interpreted as ‘light-seeking behavior’ and termed ‘dark photokinesis’[13, 46]. Zebrafish larvae lacking the eyes and pineal organ still react to a sudden loss of illumination with an elevated locomotor activity and an undirected light-seeking behavior [13]. After a few minutes of continued darkness, zebrafish will subsequently decrease their amount of swimming, resulting in less distance moved during the remaining darkness time [13, 47].

We first evaluated the responses of free-swimming *tmt-opsin1b* mutant juveniles (20-22 dpf) to sudden light changes. Mutant vs. wildtype medaka were subjected to 30min each of white light vs. dark intervals (repeated three times, spectra: S3E Fig, S4A Fig), while the movement of the larvae was tracked automatically and evaluated using the Noldus EthoVision XT software. Both *tmt-opsin1b* mutant and wildtype fish changed their swimming distance depending on the light / dark condition (Fig 2A,B, S2 Data), and there was a significant difference between both during the dark phases (Fig 2A,B, S2 Data). As there are more than 40 Opsins present in teleosts, we next wondered if this phenotype became stronger under light conditions that are more restricted to the maximal sensitivity of TMT-opsins. TMT-Opsins are functional photoreceptors with a maximal sensitivity in the blue light range (about 460nm [14, 32, 48, 49]). Thus, we next modified the assay by using monochromatic blue light (for spectra see S3F Fig, S4A Fig). Under these light conditions *tmt-opsin1b* mutant fish exhibited a clear increase of activity during the light and dark phases of the assay compared to wt (Fig 2C,D, S2 Data). Given this clear difference between white and blue light conditions, we next wondered if this response is specific to blue light. We assayed juvenile fish under lower intensity monochromatic blue, green and red light at as similar photon numbers as possible. Under these conditions we find that juvenile wt and *tmt-opsin1b*−/− mutant fish assayed under blue light, but not under green or red light, exhibited statistically significant differences (Fig 2E-G,H,J,L, S2 Data, for spectra see Fig 2I,K,M, S3G-I Fig). These results show that the response is specific and provides additional evidence for the blue light sensitivity of TMT-opsin1b.

**Fig 2:**
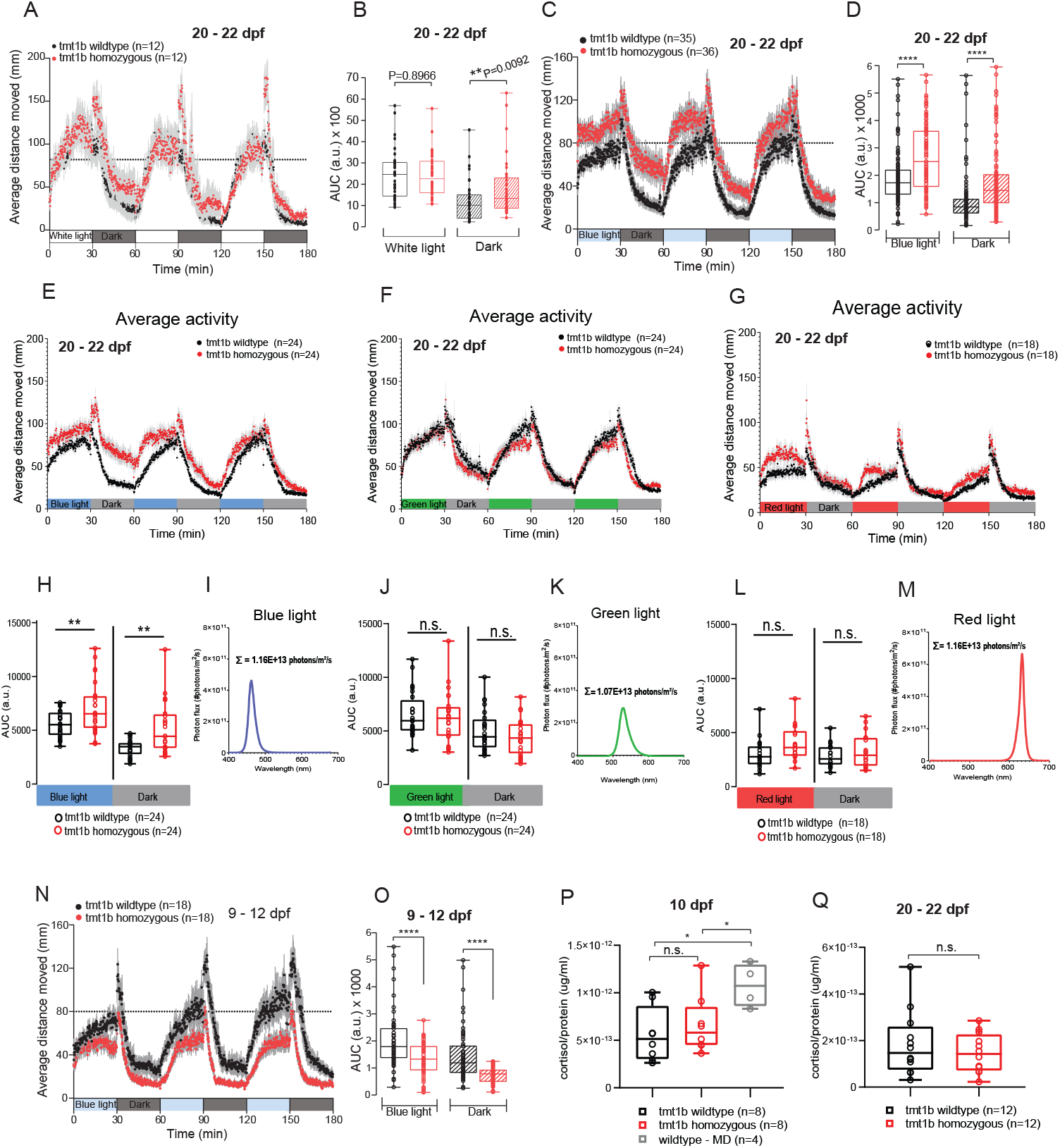
Mutations in *Ola*-*tmt-opsin1b* impact on activity levels upon light changes in a partly age-dependent manner. (**A-G, H,J,L,N,O**) Average distance moved and corresponding statistical analysis during alternating 30 minutes white light or monochromatic light/ dark intervals from juvenile fish (A-G,H,J,L) and larvae (O,P). Each data point (A,C,E-G,N) represents the mean (± s.e.m.) distance moved for the preceding 10 seconds. Colored boxes on x-axis indicate light condition. (**B**,**D,H,J,L,O**) Locomotor activity measured as area under the curve: AUC during the entire light vs. dark intervals of the trial. \*\*\*\**P* ≤ 0.0001; ** *P* ≤ 0.01. (**I,K,M**) Plotted spectra of the different light conditions used in E-G,H,J,L. The sum of photons is shown next to the respective spectrum. (**P,Q**) Cortisol levels measured as cortisol (ug/ml) over total protein levels (ug/ml) under baseline and mechanically disturbed (MD) conditions from larvae (N) and juvenile fish (O). \**P* ≤ 0.05. For all panels, (black) *tmt-opsin1b* and (red) *tmt-opsin1b* homozygous single mutant. Box plots: single bar: median, box: first and third quartile, whiskers: min and max; n.s.: no significance; a.u.: arbitrary units. Metadata: S2 Data. For further details on spectra and age-dependent differences: S3 Fig, S4 Fig.

Many behavioral and brain imaging experiments with genetically altered and/or transgenic zebrafish (as major functional vertebrate model system in neuroscience) are typically performed at early developmental stages when the fish have just started to swim freely (7dpf) or even earlier (e.g [50, 51]). As detailed in the introduction, also all analyses of non-visual photoreceptor responses in teleosts have so far only been performed at stages not older than 7dpf, and in most cases significantly younger [13, 20, 21] [22].

However, it is also obvious from multiple studies that significant improvements of sensory systems and behavior occur during subsequent development (e.g. [52, 53]).

We thus next tested how our results with juvenile fish compare to those at stages just post-hatching. The result was strikingly different. While there was also clearly a difference compared to wildtype, instead of swimming more, mutant young larvae (9-12 dpf) swam less than their wildtype siblings (Fig 2N,O, S2 Data). This behavioral difference changed gradually during the days following post-hatching (S4B-I Fig, S8 Data), emphasizing that any molecular or cellular mechanism identified to control behavior has to be seen in connection with the developmental age of the tested organism.

Analyses of the “fast dark photokinesis” response rate (first two minutes after light cessation) revealed no difference in the amount of the response between *tmt-opsin1b* mutant and wildtype of either age, when the overall lowered or heightened mutant baseline level is accounted for (S4J,K Fig, S8 Data).

Finally, in order to test if the observed elevated locomotion levels are caused by generally altered stress or anxiety levels, we measured total cortisol levels.

We observed no difference between mutant and wildtype siblings at neither young larval nor juvenile stages (Fig 2P,Q, S2 Data), while our positive control, larvae exposed to mechanical disturbances (MD), showed a significant increase of cortisol levels (Fig 2P, S2 Data). This strongly suggests that the difference in locomotion apparent between mutants and wildtype is indeed mediated by the acute light changes involving *tmt-opsin1b*.

### *Ola-tmt-opsin1b* mutants exhibit altered, partly eye-independent, daytime activity levels

Given the relatively consistent changes over the entire period of the experiment, we next tested the free-swimming of *tmt-opsin1b* mutants across two consecutive days (blue light/dark, 16h:8h). *Tmt-opsin1b* homozygous mutant larvae and juvenile fish swam significantly more during the 1.5h period immediately after the lights went on (Fig 3A-D, S3 Data). This difference was reduced or disappeared when the light conditions remained stable over the rest of the day (Fig 3A,C, S3 Data).

**Fig 3:**
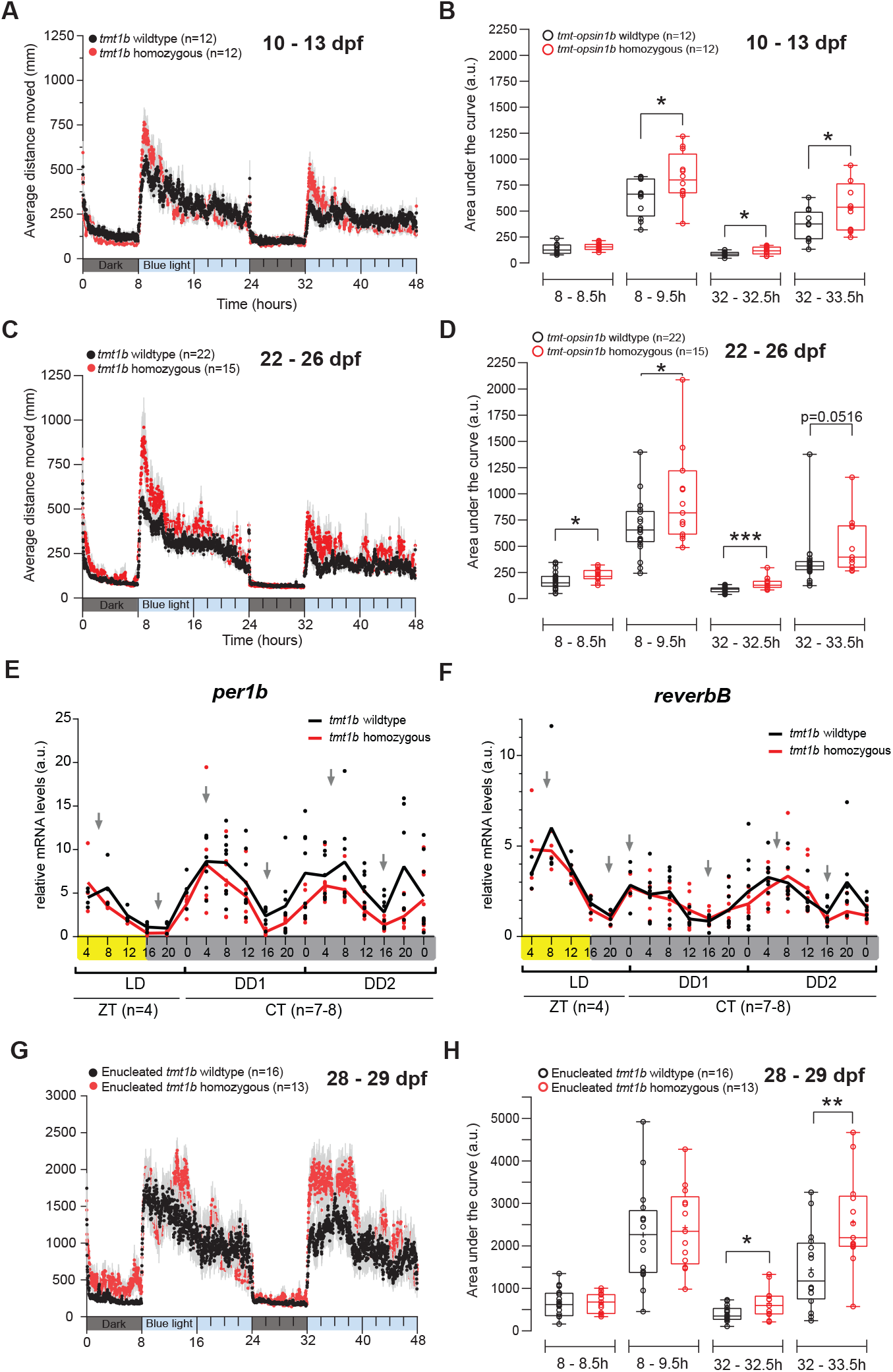
*Tmt-opsin1b* affects the amount of daytime rest, independently of eyes and the circadian core clock. (**A,C,G**) Average distance moved during 8 hours dark / 16 hours blue light periods for two consecutive days in larvae (A), juvenile fish (C) and enucleated juvenile fish (G). Each data point represents the mean (± s.e.m.) distance moved for the preceding 1 minute. Colored boxes along the x-axis represent light condition. (**B,D,H**) Locomotor activity (measured as AUC) during the first 30min and 90min after light is turned ON during the 2 days trial. (B) \**P*_(8-9.5h)_ = 0.0331; \**P*_(32-32.5h)_ = 0.0247; \**P*_(32-33.5h)_ = 0.0323; (D) \**P*_(8-8.5h)_ = 0.0330; \**P*_(8-9.5h)_ = 0.0299; \*\*\**P*_(32-32.5h)_ = 0.0002; (F) \**P*_(32-32.5h)_ = 0.0192; \*\**P*_(32-33.5h)_ = 0.0056. (**E,F**) Relative mRNA levels of per1b (E) and reverb (F) measured from whole larvae entrained to 16:8 white light/dark cycles. Yellow and gray bars: light vs dark conditions during respective sampling. Gray arrows indicate the peaks and troughs of the curve. For all panels, (black) *tmt-opsin1b* and (red) *tmt-opsin1b* homozygous single mutant. Box plots: single bar: median, box: first and third quartile, whiskers: min and max; n.s.: no significance; a.u.: arbitrary units. Metadata: S3 Data. For further details on spectra and individual, more time-zoomed movement plots: S3 Fig, S5 Fig.

Light is an important entrainment cue for the circadian oscillator. We thus next analyzed, if the mutation in *tmt-opsin1b* impacts on the phase or period length of the medaka circadian core clock. We selected two representative core circadian clock genes, *per1b* and *reverbB* [54], and tested their transcript oscillations in *tmt-opsin1b* mutant and wt larvae under light/dark (L/D) and constant dark (D/D) conditions. The timing of daily minima and maxima was indifferent between mutant and wt larvae, as was the overall cycling, clearly indicating that the *tmt-opsin1b* mutation has no impact on circadian phase or period length (Fig 3E,F, S3 Data).

Finally, we tested, if the increase in swimming upon sudden illumination would also occur in fish without eyes, by this testing for a possible functional contribution of *tmt-opsin1b* outside the eye (and pineal, as *tmt-opsin1b* is not expressed in the pineal [14]). Fish were enucleated and left to recover for 1 week before the trial onset. Importantly, the differences between *tmt-opsin1b* and their wildtype counterparts were still observable: while the trend was the same during both days, it reached statistical significance only during the second day (Fig 3G,H, S3 Data). We thus conclude that *tmt-opsin1b*, at least in part, functions outside the eyes and modulates the responses of medaka behavior in response to different environmental light changes independently of the circadian clock.

### *Ola-tmt-opsin1b* and *Ola-tmt-opsin2* mutants exhibit additive and non-additive responses to changes in environmental light

Teleost, like zebrafish and medaka, possess more than 40 Opsins. Systematic analyses in zebrafish showed that most exhibit expression in the brain [15]. This raises the question if these opsins might just function redundantly or rather in a complex, non-redundant manner. We purposely started to approach this question by investigating an additional Opsin with similar characteristics as Tmt-opsin1b. Specifically, Tmt-opsin2 is an evolutionarily conserved ETO relative of Tmt-opsin1b with highly similar spectral sensitivity and absorbance characteristics [14, 32] and expression in adjacent, possibly partly overlapping, domains in mid- and hindbrain [14] (Fig 4A-E).

**Fig 4:**
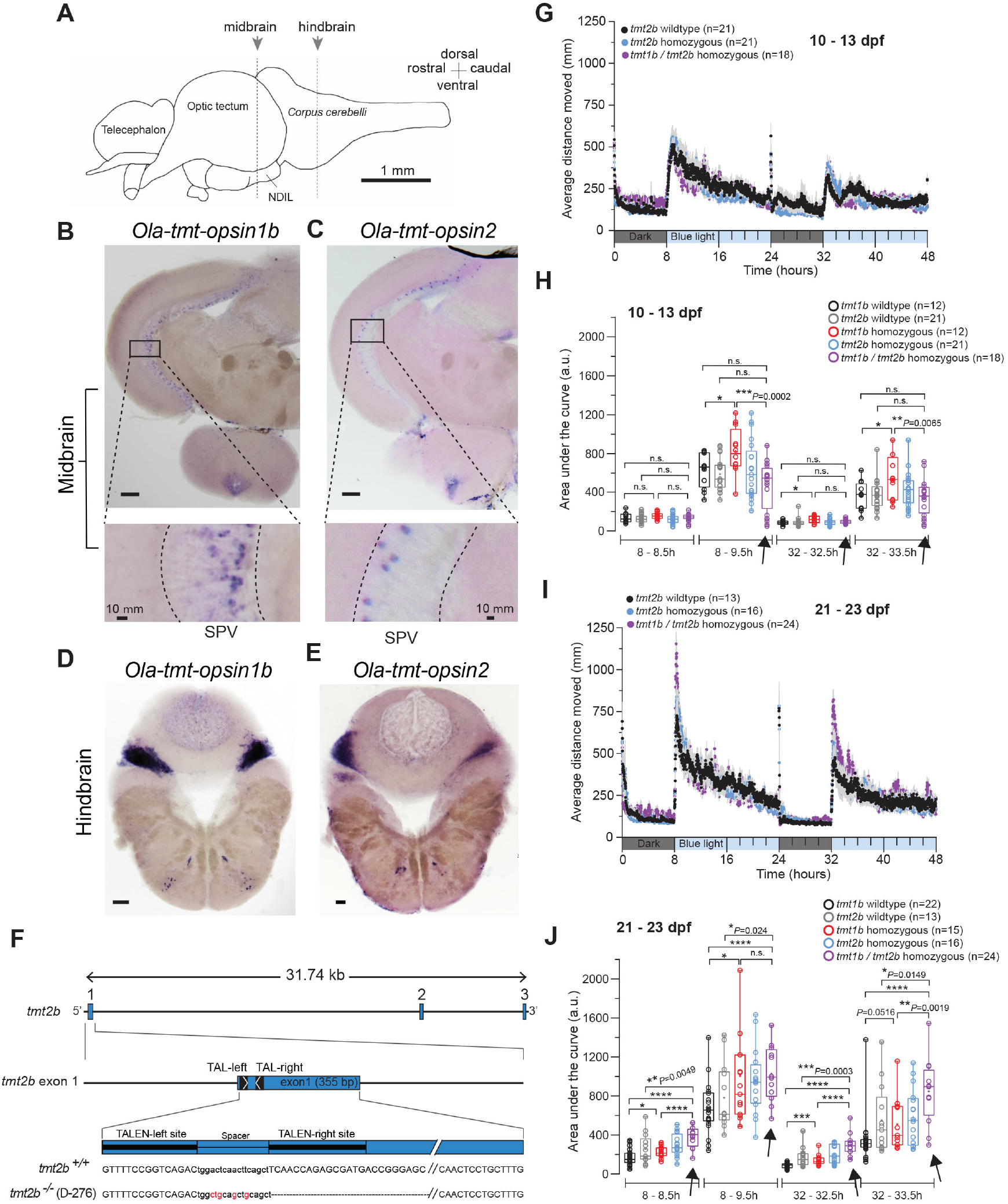
Individual and combined differential impact of *Ola-tmt-opsin1b* and *Ola-tmt-opsin2* on daytime rest levels. (**A**) Schematized lateral view of an adult medaka brain. Dashed lines: position of sections in B,C and D,E. NDIL: nucleus diffuses of lobus inferiosis (of hypothalamus). (**B**,**C**,**D**,**E**) *In situ* hybridization (ISH) of *tmt-opsin1b* (B,D) and *tmt-opsin2* (C,E) on coronal sections from Cab wildtype fish. SPV: *stratum periventriculare*. Scale bars: 100 μm. For detailed anatomical annotations see [14] (**F**) Genomic locus of the *Olatmt-opsin2* gene and corresponding mutant. TALEN binding sites: black boxes; Exons: blue boxes; Inserted/substituted nucleotides: red; deleted nucleotides: “−”. (**G**,**I**) Average distance moved profile during a 2 consecutive days trial on a 16h:8h blue-light/dark photoperiod of (G) 10-13 dpf larvae and (I) 21-23 dpf juvenile fish. Each data point represent the mean (± s.e.m.) distance moved for the preceding 1 minute. Colored boxes along the x-axis represent light condition. (**H**,**J**) Locomotor activity (assessed by area under the curve) during the first 30 minutes or 90 minutes after light is turned ON along the 2 days trial of (H) larvae and (J) juvenile fish. Arrows highlight the data from the double-mutants to be compared to the respective single mutants and wt. For all panels: \*\*\*\**P* < 0.0001. Box plots: single bar: median, box: first and third quartile, whiskers: min and max; n.s.: no significance; a.u.: arbitrary units. Metadata: S4 Data. For further details on mutant, spectra and individual, more time-zoomed movement plots: S1 Fig,S3 Fig,S5 Fig.

We thus wondered, if mutating *tmt-opsin2* would reveal redundant or synergistic functions with *tmt-opsin1b*. Using TALE nucleases, we generated a large deletion in the *tmt-opsin2* gene (Fig 4F), removing the first two transmembrane helices and resulting in a dysfunctional protein (S1E-G Fig). Larval and juvenile *tmt-opsin2* mutant and wildtype sibling fish were assessed for their responses during the light / dark phases across two days, as described above.

While under these conditions *tmt-opsin2*−/− mutants do not display significant phenotypes, unexpectedly *tmt-opsin1b/tmt-opsin2* double homozygous mutants displayed phenotypes different from the *tmt-opsin1b*−/− single mutants: while in larval stages adding the *tmt-opsin2* mutant to the *tmt-opsin1b* mutant resulted in a complementation of the *tmt-opsin1b* phenotype during the beginning of the light phases (Fig 3A,B, S3 Data, Fig 4G,H, S4 Data, for individual, high temporal resolution graphs see S5A-J Fig), the combination of these mutations lead to additive phenotypes in the same experimental test during juvenile stages (Fig 3C,D, S3 Data, Fig 4I,J, S4 Data, for individual, high temporal resolution graphs see S5A-J Fig; compare position of purple bar graphs -also indicated by arrows- to the corresponding wt and single mutant graphs in Fig 4H and J, S4 Data).

These data suggest two main conclusions. First, the fact that the *tmt-opsin2* mutation can compensate for *tmt-opsin1b* loss shows that these photoreceptors do not function “just” redundantly. It suggests that light information received by the fish is fed into a complex light processing system and the output for behavior is not simply the sum of the input of all possible light receptors. Second, the functional interaction of the different opsin-based photoreceptors has a clear age-dependent component.

### Mutations in *tmt-opsin1b* and *tmt-opsin2* induce transcriptional changes that impact on neuronal information transmission

We next wondered which molecular changes can be detected in the brain of the fish missing specific photoreceptors. It had previously been shown that differences in photoperiod cause changes on the transcript level in the rat brain, resulting in differences of neurotransmitter abundance [7]. We thus reasoned that quantitative RNA sequencing might be a possible strategy to obtain an unbiased insight into the changes that occur due to lack of *tmt-opsin1b* or *tmt-opsin2* function.

We sampled three separate brain regions (eyes, forebrain and mid-/hindbrain, Fig 5A) at ZT2-3 under a 16:8 hour white light/dark regime and sequenced stranded cDNA. It should be noted that for technical reasons part of the mid-/hindbrain sample includes a portion of the posterior hypothalamus, i.e. forebrain. The resulting sequences were mapped to the medaka genome (Ensembl version 96), reads mapping to annotated exons were quantified using edgeR (Bioconductor version 3.9) [55] . The result tables are deposited as S10 Data, S11 Data. When comparing *tmt-opsin1b* and its wildtype siblings, one differentially regulated transcript caught our particular attention, the pre-pro-hormone *sst1b* (ENSORLG00000027736: also named *sst3* or *cort*) in the mid-/hindbrain (Fig 5B, S5 Data), a member of the somatostatin/corticostatin family. We next independently confirmed the quantitative RNAseq results using qPCR (Fig 5C, black vs. red boxes for mid- and hindbrain, S5 Data). In this set of experiments, we separated the midbrain from the hindbrain tissue in order to obtain a more differentiated picture of the regulation. Given the differential effects of *tmt-opsin1b/tmt-opsin2* double mutants on behavior, we next wondered how the presence of both mutations would affect *sst1b* regulation. Adding the *tmt-opsin2* mutation compensated the downregulation of *sst1b* levels present in the *tmt-opsin1b* single mutant (Fig 5C, purple boxes compared with others for mid- and hindbrain, S5 Data). In an analogous approach, we identified the voltage-gated sodium-channel subunit *scn12aa* as significantly regulated by the *tmt-opsin2* mutation (Fig 5D, S5 Data). Again, while clearly visible in the single mutant (Fig 5E, grey vs. blue boxes for mid- and hindbrain, S5 Data), the effect was compensated for in the *tmt-opsin1b/tmt-opsin2* double mutants (Fig 5E, purple vs. blue boxes for mid- and hindbrain, S5 Data).

**Fig 5:**
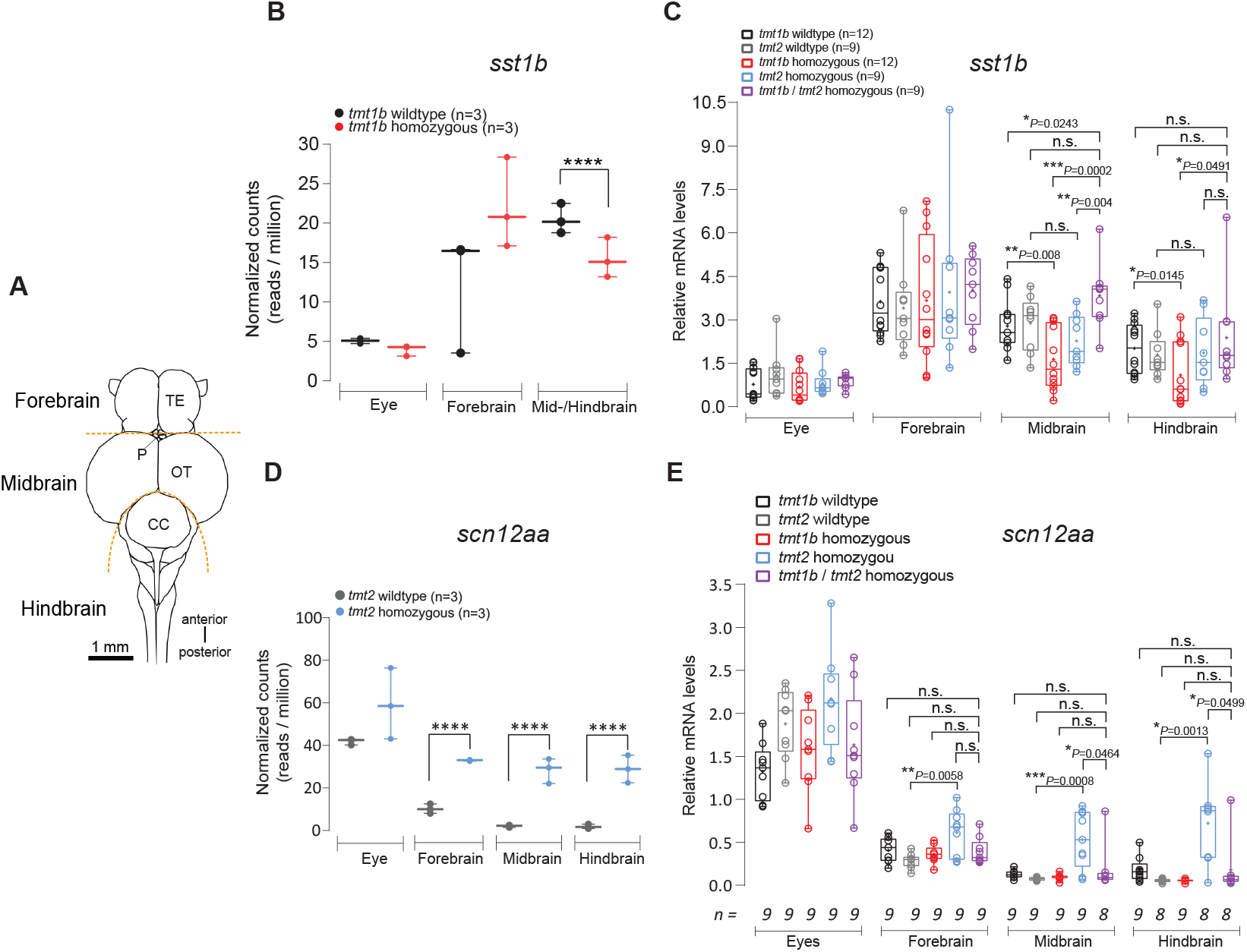
*Ola-tmt-opsin1b,2* double mutants are compensated for differential transcript changes present in *Ola-tmt-opsin1b* and *Ola-tmt-opsin2* single mutants. **(A)** Schematic drawing of the adult medaka brain in dorsal view. Dotted lines indicate the dissection boundaries (TE: telencephalon; P: pineal organ; OT: optic tectum; CC: corpus cerebelli). Note: part of the posterior hypothalamus is dissected with the midbrain, as it is attached to its ventral part and technically unfeasible to reliably separate. **(B)** Normalized transcript read counts (**** adjusted P-value < 0.001) for *sst1b*. **(C)** mRNA transcript levels for sst1b in brain areas and eyes. Decreased sst1b-levels are unique to *tmt-opsin1b*−/− mutants and are compensated by the *tmt-opsin2*−/− mutant. **(D)** Normalized transcript read counts (**** adjusted P-value < 0.001) for *scn12aa*. **(E)** mRNA transcript levels for *scn12aa* in brain areas and eyes. Increased levels of *scn12aa* are unique to *tmt-opsin2*−/− mutants, and compensated for when both *tmt-opsin1b* and *tmt-opsin2* are jointly mutated. Box plots: single bar: median, box: first and third quartile, whiskers: min and max, each displayed data point corresponds to one individual biological replicate. a.u.: arbitrary units. Metadata: S5 Data.

These molecular data further support the notion that the loss-of-function of non-visual Opsins does not necessarily lead to a simple summation effect originating from the single mutants. Again, this strongly corroborates the notion of non-redundant, complex light information processing, which can modulate the neuronal function on the level of neuropeptides (*tmt-opsin1b* regulating *sst1b*) and voltage-gated channels (*tmt-opsin2* regulating *scn12aa*).

### The regulation of *sst1b* via *tmt-opsin1b* is non-cell autonomous

In order to gain deeper insight on the connection between *sst1b* expression changes in the mid-/hindbrain, *tmt-opsin1b* and responses to light, we next analyzed the spatial expression of *sst1b*. In the midbrain, *sst1b* is expressed in several highly specific clusters of cells, none of which overlap with *tmt-opsin1b* (compare Fig 6A and Fig 4B, for detailed anatomical annotation of *sst1b* expression see S6 Fig, of *tmt-opsin1b* expression see [14]). This is particularly obvious for the tectum, in which *sst1b*+ cells are consistently located more dorsally within the *stratum periventriculare* (SPV) than *tmt-opsin1b+* cells, and close to or possibly overlapping with *tmt-opsin2+* cells (compare Fig 6A with Fig 4B, anatomical annotation: [14], S6 Fig). Similar in the reticular formation of the hindbrain, likely all of the *sst1b+* cells are separate from the *tmt-opsin1b+* cells (Fig 6B,4D, anatomical annotation: [14], S6 Fig). We thus conclude, that the changes of *sst1b* transcript levels are rather indirectly mediated by *tmt-opsin1b+* cells upstream of *sst1b*-expressing cells.

**Fig 6:**
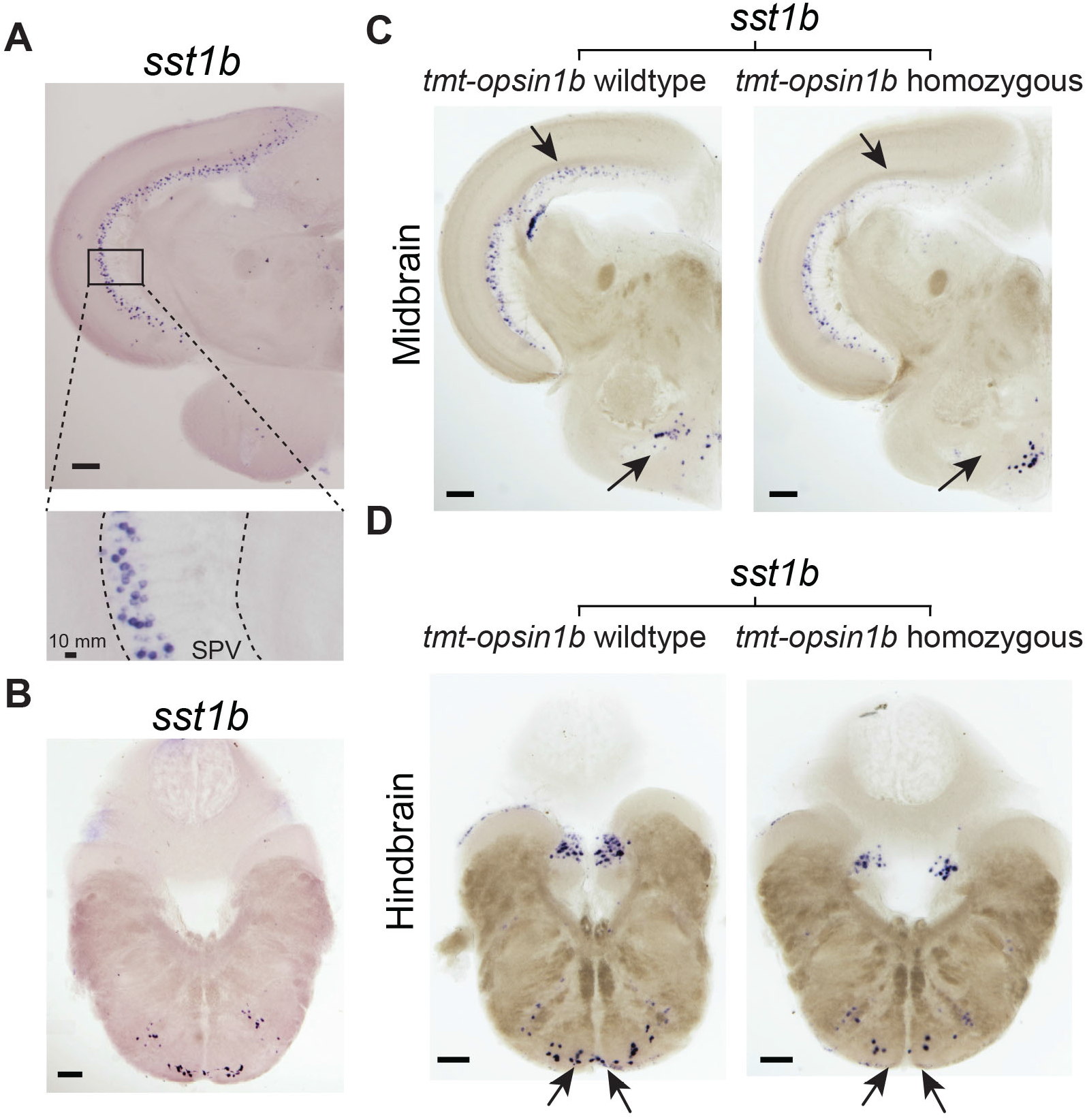
*sst1b* levels are reduced in specific cells in the tectum, posterior hypothalamus and hindbrain in *tmt-opsin1b*−/− medaka brains. *In situ* hybridization for *sst1b* performed on coronal sections of midbrain **(A)** and hindbrain **(B)** from Cab wildtype fish. Magnification of boxed areas of the optic tectum are displayed below. SPV: stratum periventriculare. Note that *sst1b* identify a distinct tectal population of dorsal cells and on the ventral area of the hindbrain. **(C,D)** Comparison of *sst1b* expression in wildtype vs. *tmt-opsin1b* mutant sibling coronal brain sections reveals specific reduction (denoted by arrows) in interneurons of the tectum and reticular formation, and in the posterior hypothalamus. Scale bars: 100um. For further details on anatomical annotations, co-localization of *sst1b* with *gad2,* as well as Scn12aa orthology relationships and localization: S6 Fig, S7 Fig, S8 Fig, S9 Fig.

We next analyzed where the regulation of *sst1b* transcripts occur. We performed *in situ* hybridizations on wildtype and *tmt-opsin1b* mutant adult fish brains for *sst1b* with specific attention to treating wildtype and mutant samples identically. We observed a specific reduction of *sst1b*+ transcript levels in the cells of the SPV layer in the optic tectum, the absence of *sst1b* staining in the dorsal periventricular hypothalamic zone surrounding the lateral recess and in cells medial to the nucleus glomerulosus in the medial preglomerular nucleus (arrows Fig 6C). We also observed a reduction of *sst1b*+ cells in the intermediate reticular formation of the hindbrain (arrows Fig 6D). In order to independently verify the specificity of the effects, we cloned *gad2,* whose ortholog is co-expressed with the zebrafish *sst1b* ortholog in neurons, based on scRNA analyses [56]. We confirmed co-expression of *sst1b* with *gad2* in medaka tectal neurons (S7 Fig), and also did not find any visible alteration in the expression of *gad2* in *tmt-opsin1b* mutants vs. wt brains (S7 Fig). Taken together, these *in situ* expression analyses confirm a likely specific effect of *tmt-opsin1b* on *sst1b* transcripts. They further suggest an indirect regulation of *sst1b* transcripts by *tmt-opsin1b.* We additionally identified that the tectal *sst1b+* cells are GABA-ergic (by the expression of *gad2*), and hence likely convey inhibitory input to the tectal circuitry.

We likewise further analyzed *scn12aa* brain expression. *Medaka* Scn12aa is the common ortholog of the two separate amniote Scn10a and Scn5a groups (S8 Fig, S9 Data).

Expression analyses of both mRNA (by *in situ* hybridization, S9A,B Fig) and antibody staining (using a commercially available antibody with broad species reactivity generated against human Scn5a, S9C-H Fig) suggest a broad, possibly ubiquitous brain expression. A specific set of neurons regulated by *tmt-opsin2* was not obviously detectable. It is hence plausible that the effect of *tmt-opsin2* is present in multiple neuronal populations and not restricted to few specific circuits.

### Photoperiod regulates *sst1b* transcript levels via a *tmt-opsin1b* dependent mechanism

We finally aimed to further investigate the possible light dependency of the observed transcript regulation. Given that *sst1b* exhibits the more specific expression, we focused on this transcript. Specifically, we tested, if changes in the illumination regime impact on *sst1b* transcript levels in a TMT-Opsin1b dependent manner. We chose a ‘photoperiod’-type light regime based on three reasons: First, any changes in transcription need sufficient time to occur. Second, we wanted to test a light regime that has obvious natural relevance to medaka [30], and third, *pre-pro-somatostatin* expression is regulated in response to exposure to short- and long-day photoperiods in adult rats [7].

We exposed wildtype fish to two different white light regimes (Fig 7A,B). All fish were initially raised under a 16h (light):8h (dark) white light regime (Fig 7B). One cohort remained exposed to this 16h:8h LD cycle (long day), while the other group was transferred to an 8h:16h LD cycle (short day). After one week eyes and brains were dissected at ZT 8 (long-day) and ZT4 (short-day) (blue arrowhead Fig 7B) and analyzed by qPCR.

**Fig 7:**
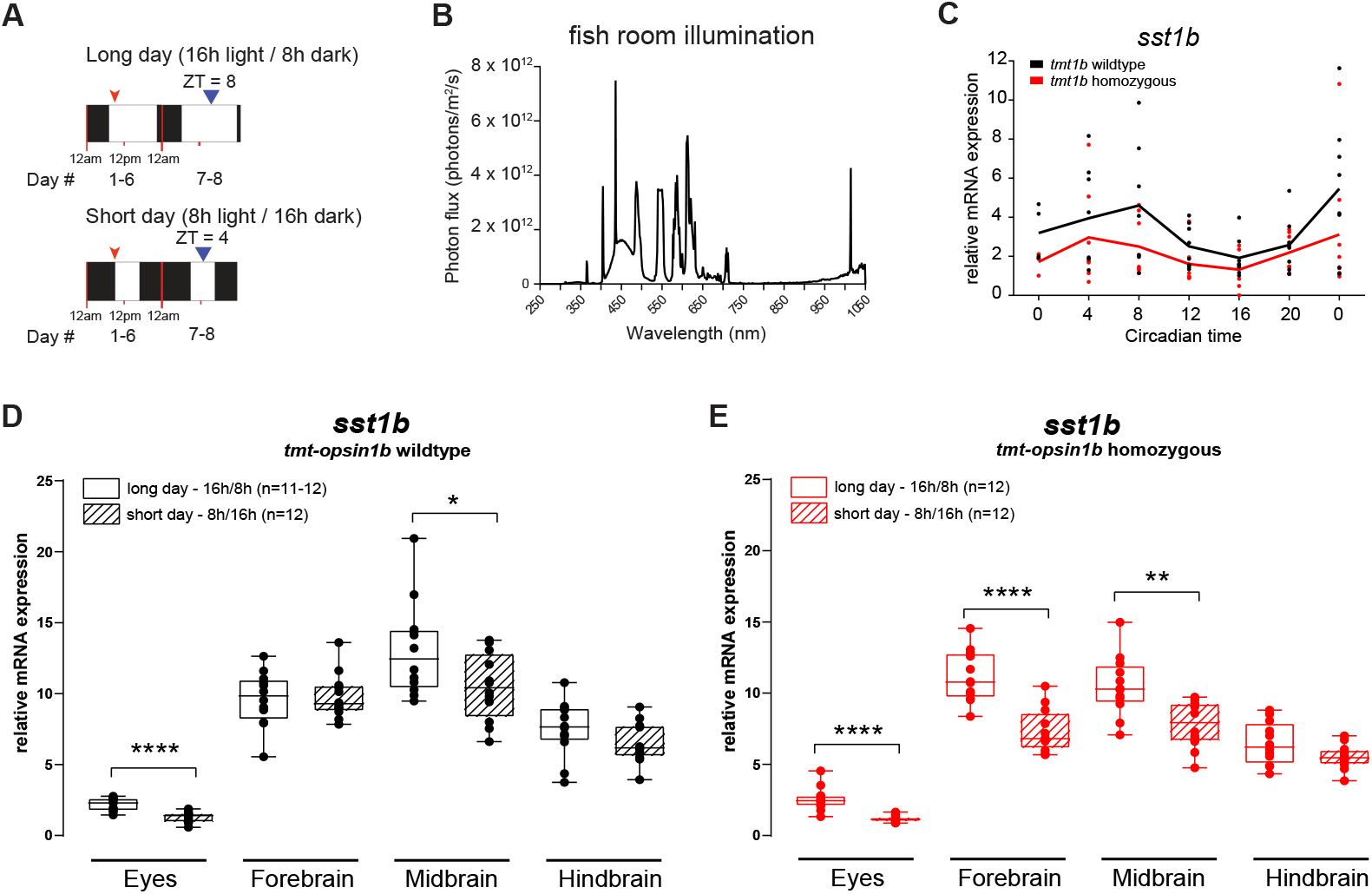
*tmt-opsin1b* sensitizes *sst1b* regulation to changes in photoperiod. (A) Schematic representation of the two different photoperiods assigned to different cohorts of *tmt-opsin1b* wildtype and *tmt-opsin1b* mutant adult fish. Black boxes: dark period; White boxes: white light period. Red arrowheads: feeding time; Blue arrowheads: time when fish were sacrificed and the brain dissections made. (**B**) Light spectrum of the light used in (A). (**C**) Relative mRNA expression of *sst1b* in larvae sampled during constant darkness after entraining with 16:8 hour white light/dark cycle at 27°C. (**D,E**) mRNA *sst1b* transcript expression levels in *tmt-opsin1b* wildtype (D), *tmt-opsin1b* mutant siblings (E). \*\*\*\**P* ≤ 0.0001, \*\**P* ≤ 0.01, \**P* ≤ 0.05. Box plots: single bar: median, box: first and third quartile, whiskers: min and max. Each data point represents one biological replicate. Metadata: S6 Data. For further details on spectra: S3 Fig.

As *sst1b* is circadianly regulated with no phase difference between *tmt-opsin1b* mutants and wt fish (Fig 7C, S6 Data), we timed our brain part sampling to the middle of the respective light phases, an established sample timing strategy for photoperiod analyses in medakafish [57].

A comparison of *sst1b* transcript levels at long vs. short day light regimes revealed significantly lower transcript level for the short day cohort, mimicking the *tmt-opsin1b* mutation (compare Fig 7D vs. Fig 7E, S6 Data and red vs. black boxes Fig 5B,C, S5 Data). Comparing the transcript levels of *sst1b* under long day vs. short day photoperiods in the *tmt-opsin1b* mutants revealed that this mutation leads to even stronger changes in *sst1b* transcript levels (Fig 7E, S6 Data). This is most apparent for the forebrain, where there are no changes between long- and short-day conditions in wildtype (Fig 7D, S6 Data), but a strong downregulation of *sst1b* under short-day in the mutant (Fig 7E, S6 Data). This shows that the regulation of *sst1b* by light depends on *tmt-opsin1b,* but – similar to the behavioral changes in the avoidance assay– suggests that *tmt-opsin1b* normally prevents specific neurophysiology changes to occur in response to environmental light changes, a “stabilizing function” that is perturbed in *tmt-opsin1b* mutants.

## Discussion

The large number of *Opsin* genes uncovered over the past years in vertebrates, particularly in teleosts, is puzzling. Even more since many of these *Opsins* have been shown to encode functional light receptors and are expressed at various places beyond the rods and cones (e.g. [9-12, 14-17]). Here we have started to functionally tackle this diversity by investigating loss-of-function mutants of two non-visual Opsins of the Encephalopsin/TMT-Opsin family in medaka.

The results are multifaceted, complex and provide food for thought from multiple perspectives.

While first focusing on *tmt-opsin1b*, we show that this Opsin modulates fish swimming behavior in a light-dependent manner in several assays. Interestingly, in most cases do *tmt-opsin1b* mutant fish display enhanced behavioral responses. At present we can only speculate about the contribution of the different *tmt-opsin1b+* neurons in the response circuitry. While the contribution of several *tmt-opsin1b+* cells is unclear, it is highly likely that tectal cells are involved. This may suggest the following circuitry: *Tmt-opsin1b+* tectal cells are positive for ChAT [14]. These cells connect –possibly monosynaptically (given the close vicinity and branching of ChAT tectal neurons-see [14])– to neurons marked by *sst1b/gad2*. We showed that Tmt-opsin1b affects the production of *sst1b*. A member of the somatostatin/corticostatin family in the mouse (SST, active form SST-14) has been shown to itself suppress excitatory inputs to downstream interneurons, resulting in enhanced visual gain and orientation selectivity [58]. It might hence be plausible to assume that tectal *sst1b* also exert suppressive effects. In *tmt-opsin1b* mutants *sst1b* is downregulated in tectal interneurons, which would reduce such a suppressive effect and result in enhanced behavioral responses. Alternatively, we showed that these *sst1b+* neurons also express *gad2.* It might thus also be imaginable that GABA-release could be modulated by Tmt-opsin1 in these neurons. Thus, focusing on the tectal circuitry, there are at least two possibilities how loss of *tmt-opsin1b* results in increased activity. However, future work on specific neuronal circuits will be required to really understand the role of the different *tmt-opsin1b* expressing neurons.

We also showed that the response differences present in *tmt-opsin1b* versus wt fish, as well as the effects of the *tmt-opsin1b/tmt-opsin2* double mutants are strongly age-dependent. As the expression domains of *tmt-opsin1b* nor *tmt-opsin2* do not observably change between young larvae and adults [14], we think that the most plausible explanation for the age-dependency is the still on-going differentiation of the nervous system during young larval stages. Ontogenetic changes in behavior are very common and for instance well documented for the visual processing and social behavior in medaka fish [53, 59]. While the cellular and molecular causes still remain to be determined, it is clear that the phenomenon is common among vertebrates and even extends to humans across lifespan, where it has been shown to correlate with changes in glutamatergic and GABAergic systems [60, 61]. Especially the latter is interesting in the context of our analyses, as the *sst1b+* neurons downstream of *tmt-opsin1b* function in the tectum are GABAergic and might hence be a possible subject to ontogenetic changes. Behavioral changes across life span are not just an important topic to better comprehend the extent of natural behavioral plasticity. It is also relevant to understand behavioral adaptations to natural environments, as ontogenetic changes are often intertwined with habitat shifts to balance the different risks and needs of individuals at different ages, often correlated with size differences [62, 63].

Another level of molecular modulation we uncover is the regulation of *sst1b* by photoperiod. Short-day photoperiods result in levels of *sst1b* in mid- and hindbrain that are equal to those of the *tmt-opsin1b* mutants. However, in the mutants these levels become reduced even further, and *sst1b* reductions also occurs strongly in the forebrain, where it is unaffected in wildtype under long-vs. short-day regimes. These data suggest that *tmt-opsin1b* normally prevents molecular alterations to happen when environmental conditions change.

Interestingly, different members of the *somatostatin/corticostatin* family are frequently connected to changes in light parameters. A change of the preprohormone transcript and peptide levels of Somatostatin1 has been reported for long vs. short photoperiods in adult rats [7], suggesting a possible evolutionary conservation of this regulatory relationship. Similar to the changes we observed for *sst1b* in medaka, SST1 amounts increase when the days are longer. In the case of the rat, this is correlated with a partial decrease of the enzyme *tyrosine hydroxylase* and the number of dopaminergic neurons, as well as an alteration in stress behavior[7]. However, in addition to the changes in the forebrain (posterior hypothalamus), as they were reported for rat, we find *sst1b* to be significantly regulated in mid- and hindbrain cells. It could be that this also extends to rat and certainly suggests, that brain regions outside the hypothalamus should be analyzed, in order to understand which effects photoperiod differences might exert on mammals. It will then be a question for the future to sort out, which molecular changes are related to which behavioral changes.

Furthermore, *sst1.1* in zebrafish has been implicated in the regulation of behavioral search strategies upon sudden loss of illumination, a behavior that also depends on melanopsin-type light receptor *opn4a.* However, it remains to be clarified if *sst1.1* functions downstream or in parallel pathways to *opn4a.* It might be that different non-visual opsins relay different light information via different *somatostatin/corticostatin* members.

Finally, one of our main aims was also to obtain first insight into the functional interaction of different photoreceptors in the organism. Unexpectedly for us, both on molecular and behavioral levels, the simultaneous loss of two *tmt-opsins* can compensate for the loss of each of them individually, suggesting that they act at least in part in opposite directions on the same downstream circuitry. In this respect the expression of *tmt-opsin1b, tmt-opsin2* and *sst1b* in the tectal region is again particularly interesting. *Tmt-opsin1b* and *tmt-opsin2* tectal expression is non-overlapping, both delimiting two distinct cellular populations, within two unique layers, while *sst1b* either overlaps or is directly adjacent to *tmt-opsin2* positive cells. This spatial expression raises the possibility that both *tmt-opsins* (and *sst1b*) might together contribute to the local neuronal circuit outlined above. Future work is needed to not only further delineate these interactions, but to also better understand their significant complexity in the context of natural light habitats and adaptations.

## Materials and Methods

### Medaka fish rearing and ethics statement

All animal research and husbandry was conducted according to Austrian and European guidelines for animal research (fish maintenance and care approved under: BMWFW-66.006/0012-WF/II/3b/2014, experiments approved under: BMWFW-66.006/0003-WF/V/3b/2016, which is cross-checked by: Geschäftsstelle der Kommission für Tierversuchsangelegenheiten gemäß § 36 TVG 2012 p. A. Veterinärmedizinische Universität Wien, A-1210 Wien, Veterinärplatz 1, Austria, before being issued by the BMWFW). Medaka fish (*Oryzias latipes*) strains were kept in a constant recirculating system at approximately 26-28°C in a 16h light / 8h dark cycle and bred using standard protocols [33]. Collected embryos were kept at 28°C until hatching in Embryo Rearing Medium (ERM; 0.1% (w/v) NaCl, 0.003% (w/v) KCl, 0.004% (w/v) CaCl_2_ × 2 H_2_O, 0.016 % (w/v) MgCl_2_ × 6 H_2_O, 0.017 mM HEPES and 0.0001% (w/v) methylene blue). Mutant lines were generated in and outcrossed at least four time to the Cab wildtype background. (*tmt-opsin1b* Δ+23 was outcrossed 13 times.) Where available, different mutant alleles were used interchangeably, thereby reducing the probability of effects induced by off-target mutations. Mutant versus sibling wildtype batches and batches from heterozygous crosses were position in random, alternating position relative to the respective light sources while grown prior to the assays. Parental fish grew in as similar fish room light conditions as possible.

### Design and construction of *tmt*-TALENs

Transcription activator-like effector nucleases (TALENs) targeting the first exon of medaka TMT-opsin 1b gene (Ensembl gene ENSORLG00000013181) and TMT-opsin2 (Ensembl gene ENSORLG00000012534) were designed using the TALEN targeter prediction tool (https://tale-nt.cac.cornell.edu/; TAL Effector Nucleotide Targeter v2.0) according to the following parameters: equal left and right repeat variable diresidue (RVD) lengths, spacer length of 15–20 bp, NN for G recognition, only T in the upstream base and the presence of a unique restriction site in the spacer region. TAL effector modules were assembled and cloned into the array plasmids pFUS using the Golden Gate TALEN and TAL Effector kit (Addgene, Cambridge, USA) according to validated procedure (Cermak et al., 2011). The RVD sequence of the left *tmt-opsin1b*-TALEN was HD NN NG NG HD HD HD NI NI HD NN HD NN NI NN HD HD NG binding to the genomic sequence 5’-CGTTCCCAACGCGAGCCT-3’ and the RVD sequence of the right *tmt-opsin1b*-TALEN was NN HD NG HD HD NN HD NG NN HD NN NG HD HD HD HD NN NG binding to the genomic sequence 5’-GCTCCGCTGCGTCCCCGT-3’ on the opposite strand featuring a unique BssHII restriction site in the 17 bp spacer region. The RVD sequence of the left *tmt-opsin2*-TALEN was NN NG NG NG NG HD HD NN NN NG HD NI NN NI HD binding to the genomic sequence 5’-GTTTTCCGGTCAGAC-3’ and the RVD sequence of the right *tmt-opsin2*-TALEN was NI NG HD NN HD NG HD NG NN NN NG NG NN NI binding to the 5’-CATCGCTCTGGTTGA-3’ genomic sequence. The final pCS2+ backbone vectors, containing the homodimeric FokI nuclease domains, were used as previously[64]. All final TALENs pCS2+ expression vectors were sequence-verified using 5’-TTGGCGTCGGCAAACAGTGG-3’ forward and 5’-GGCGACGAGGTGGTCGTTGG-3’ reverse primers.

### Generation of capped TALEN mRNA and microinjection

Full-length TALEN plasmids were digested with KpnI, gel purified (Gel Extraction kit, Qiagen, Netherlands), and *in vitro* transcripted (Sp6 mMESSAGE mMACHINE kit, Thermo Fisher Scientifics, USA), followed by a purification using the RNeasy Mini kit (Qiagen). The yield was estimated by NanoDrop (Thermo Fisher Scientific) and diluted to the final concentration for microinjection. Cab strain zygotes were microinjected with a mix containing 5 or 50 ng/μl of each transcribed TALEN mRNA and 0.6% tetramethylrhodamine (TRITC; Thermo Fisher Scientific) in nuclease-free water.

### Genotyping of *tmt-opsin1b* and *tmt-opsin2*-TALEN mutated fish

Genomic DNA from hatched larvae or caudal fin biopsies were retrieved with lysis buffer (0.1% SDS; 100 mM Tris/HCl, pH 8.5; 5 mM EDTA; 200 mM NaCl; 0.01 mg/ml Proteinase K) at 60°C overnight and 1 μl of a 1:20 dilution of the lysate used in a PCR reaction (HotStarTaq, Qiagen). The sequence flanking the *tmt-opsin1b* TALEN binding sites was amplified using the forward 5’-GGGACTTTCTTTGCGCTTTA-3’ and the reverse 5’-CAGGTCAGAGCGGATCTCAT-3’ primers. 10 μl of the reaction was directly used for restriction digest using 4 units of BssHII enzyme (New England Biolabs, USA) in a 20 μl reaction. Genotyping of the *tmt-opsin2* locus was made with the forward 5’-CGGTGAGCGATGTGACTG-3’ and the reverse 5’-GGGAGATCTTTGTCCAGGTG-3’ primers. Mutation caused by TALENs was assessed by analyzing the band sizes of the restriction digest product on a 2% agarose gel. Undigested bands were gel extracted and subcloned into pJet2-1 using the Clone JET PCR Cloning kit (Thermo Fisher Scientific) and sequence-verified.

### Immunohistochemistry

#### Without clearing

Adult fish (> 2 months-old) were anesthetized in fish water containing 0.2% tricaine, decapitated, the brain dissected, fixed in ice-cold 4% PFA overnight and subsequently stored in 100% methanol for at least 1 week before use. After rehydration in successive steps of PBT (PBS with 1% Triton X-100), the dissected brains were fixed in 4% paraformaldehyde in PBT for 2 hours, embedded in 3% agarose and cut into slices of 100 μm thickness on a tissue vibratome, followed by 4 hours of blocking in 5% sheep serum at room temperature. Slices were then incubated for 3 days at 4°C with a polyclonal rabbit anti-Ola-TMT-1b antibody (1:250 in 1% sheep serum). After extensive PBT washes, sections were treated with the secondary antibody goat anti-rabbit Alexa Fluor 488 (Thermo Fischer Scientific) in a 1:500 dilution in 1% sheep serum supplemented with 1:10000 of 4’,6-diamidino-2-phenylindole (DAPI), during 2 days at 4°C. Following PBT washes, slices were mounted in proper mounting medium and pictures were taken on a Zeiss LSM700 confocal scanning microscope.

#### With clearing

The protocol was adapted from [65] with the following details: adult fish (> 2 months-old) were anesthetized in fish water containing 0.2% tricaine, decapitated, brain dissected, fixed in ice-cold 4% PFA overnight. After washes in 1x PBS at room temperature (RT), the brains were incubated in pre-chilled acetone O/N at −20°C, rehydrated in 1x PBS at room temperature and digested with 10ug/ml of Proteinase K (Merck #1245680100) for 10-12 min at RT, followed by 2x glycine washes (2mg/ml) (Roth #3908.3). Brains were washed in 1x PBS, immersed in Solution 1.1 (3 hrs, at 37°C) under gentle shaking, subsequently washed in 1x PBS O/N, embedded in 3% agarose and sliced (100 μm thickness) using a vibrating blade microtome (Leica VT1000S, Leica Biosystems, Germany). The slices were then blocked for 1-2 hrs in 10% sheep serum / 1x PBS, followed by primary antibody incubation in 10% sheep serum at 4°C for 3 days (Anti-SCN5A antibody produced in rabbit (Sigma SAB2107930) 1:1000. After extensive 1x PBS washes, sections were incubated in a goat anti-rabbit Cy3 antibody (Thermo Fischer Scientific) (1:500 dilution) in 10% sheep serum, 1:10000 of DAPI were added, and incubated for 3 days at 4°C. Following PBS washes, slices were mounted in proper mounting medium and pictures were taken on a Zeiss LSM700 confocal scanning microscope.

### Avoidance assay

The behavioral assay was essentially performed as described before [40]. A single medaka larva (9 - 12 dpf; unfed) or juvenile fish (20 - 22dpf; fed) was placed in a custom-built acrylic glass chamber (18 × 14 × 305 mm) filled with 1x ERM medium (1 cm high), having a transparent bottom and opaque upward walls. Five chambers lying parallel to each other were placed on a horizontal facing computer screen, allowing for the simultaneous recording of 5 free-swimming animals in the same behavioral trial. Stimuli consisted of moving black dots (except when testing for contrast, where different abstract grey values were used) on a white background travelling at a constant speed of 13.5 mm/sec in the same direction in every trial. Each behavior trial consisted of 7 blocks of 6 dots each, each block characterized by a unique dot size, presented in a size-ascending or in a pseudo-random order. Stimuli were generated on the computer screen using custom-written programs (Ubuntu). Two consecutive blocks were presented with a 19 sec interval, during which no stimulus was displayed. The size of the dots was characterized by its diameter (in pixels) and were calculated as degrees of the larva visual field, as described before[40]. A industrial camera (acA1300-30gm GigE; Basler AG, Germany), positioned 40 cm above the computer screen, allowed the visualization of the fish-dot interactions, recording videos at a frame rate of 33 frames per second for offline analysis. The fish were allowed to accommodate to the chamber for 5 minutes before the beginning of the trial, which is longer than in previous similar assays in zebrafish [40] and *Xenopus* [37], and by this likely ensuring a robust standardization across different experiments. The experiments were performed during the natural light part of the fish light/dark cycle, between ZT2 and ZT5 and a single animal was used only in one behavior trial to avoid possible habituation (e.g. [66] or learning. Each fish-dot interaction was scored either as an approach or an avoidance when the fish swam toward or away from a moving dot, respectively [40]. A neutral interaction was scored when the dot entering the fish visual field cause no change in the initial swimming pattern of the animal or if the fish remained motionless. Approach behaviors consisted of swimming towards the dot, displaying a distinct attraction and predation behavior[66], whereas avoidance behaviors were faster than approaches, characterized by rapid swimming bouts away from the direction of the dot’s movement. All video analysis were made prior to genotyping of the fish to shield the identity of the subjects from the observer. An avoidance index [(A.I. = (number avoidances - number approaches) / (number total of dots in a block)] was used as a quantitative behavioral readout. All behavioral assays were made at a constant room temperature of 26°C, with the room lights switched off and the behavioral setup covered by a black light-impenetrable cover cloth, this way ensuring that the computer screen would be the only illumination source. The Michelson contrast [67] ((Lmax – Lmin / (Lmax + Lmin)) was used to quantify the relative difference in luminance between the moving dot and the computer screen background. Lmax and Lmin are luminance maximum and minimum, respectively. Medaka are known to have highly acute vision, as demonstrated by previous studies on the optomotor response [53, 68], showing excellent visual performance already from hatching [68].

### Photokinesis assay

Individual larvae (9 to 12 dpf; unfed; kept at a 16h:8h light cycle at 28°C until hatching), or juvenile medaka fish (20 to 22 dpf; fed) were distributed randomly across a 6-well plate containing 10 mL of 1x ERM medium per well. A behavioral trial consisted of a behavioral paradigm that evaluates the animal swimming activity (i.e. distance moved) during alternating blocks of 30 minutes light and darkness (3 hours in total per trial)[69], at a constant temperature of 27°C. An initial acclimation phase of 5 min to darkness was used before the start of the trial. When assessing the swimming responses in a more naturalistic-like assay, a photoperiod of 16h light / 8h dark was used during 2 consecutive days, at a constant temperature of 27°C. Behavioral assessment was made between ZT3 and ZT12. DanioVision (Noldus, The Netherlands) hardware was used to track the distance moved by the animals in each trial. Larval motion was tracked at 60 frames/s over the trial. In an alternate experiment, 1min pulses of white light and darkness were presented to the larvae, during a total trial duration of 7 minutes (refers to Supplementary Fig 2A). Video data was posteriorly analyzed offline by the tracking software EthoVision XT (Noldus) to calculate the average distance moved by the animal every 10 sec of the trial. When comparing the dark photokinesis between *tmt-opsin1b* mutant and wildtype siblings (refers to S2M,N Fig), we normalized the different baseline levels by taking the average distance moved during the 5 minutes preceding darkness along the trial as a normalization factor. Each animal was used only once per trial, thus avoiding possible habituation biases. The ILT950 spectrometer (International Light Technologies Inc., USA) was used to measure the spectra and intensity of the different light sources. A light-impenetrable cloth was placed over the behavior setup, thus reassuring a lack of external light contamination.

### Diel swimming test

Care was taken to start each trial at the precise light-to-dark transition, thus minimizing any deleterious effect that the shift to a new environment might impose on the natural behavior of the fish. Since scheduled food availability has an impact on the locomotor activity of fish [70], animals were kept unfed during the two consecutive days of the experiment, thus ensuring that the change in ambient light was the central feature impacting on swimming responses. The temperature was kept constant at 27°C. The setup, tracking and off-line analysis was made as previously described for the “photokinesis assays”, with the exception that the activity of the animals was averaged every 1 minute.

### Enucleations

Enucleations were performed on 13 dpf wildtype and *tmt-opsin1b* mutant juvenile fish upon anaesthetization with 0.03% tricaine (MS-222) (Sigma, St. Louis, MO). After the surgery, fish were allowed to recover overnight in fresh rearing fish water at 26°C. After 1 week post-surgery, the fish were used for behavior testing. Handling control fish were anaesthetized for the same duration and placed afterwards in fresh fish water overnight.

### RNA extraction and sequencing

For RNA-seq experimentation, the eyes, forebrain, midbrain and hindbrain (anatomical boundaries according to [71] were dissected from age-matched wildtype and mutant *tmt-opsin1b* and *tmt-opsin2* adult (>2 months old) medaka sibling fish. One metal bead (Peqlab Biotechnologie GmbH, Germany) and 350 μl RLT buffer (Qiagen) including 1% β-mercaptoethanol was added to the frozen tissue parts, followed by homogenization for 3 minutes at 30 Hz using the Qiagen tissue lyser. Total RNA was extracted using the RNeasy Mini kit (Qiagen) according to manufacturer’s protocol. Three independent biological replicates, each made of 3 individual fish (total of 9 fish per genotype) were used. For *tmt-opsin1b*, 1 fish per independent mutation was used. The quality of total RNA was checked using the Agilent RNA 6000 Nano kit (Agilent, USA) and then enriched for poly(A)+ RNA using the Dynabeads mRNA Purification Kit (Thermo Fisher Scientific). We used the SuperScript VILO cDNA Synthesis Kit (Thermo Fisher Scientific) to generate strand-specific cDNA that was further sequenced on the Illumina Hiseq2500 platform by the VBCF NGS Unit (www.vbcf.ac.at) as 100 base single-end reads, resulting on a 15-61 million reads, on average, per biological replicate.

### Differential gene expression

Sequences from each sample were mapped against the assembled chromosomes of the medaka genome (Ensembl version 96) using the read mapper NextGenMap [72]. After filtering for duplicates and low quality base, the stand-alone *featureCounts* tool [73] was used to count mapped reads per each transcript in each sample. The Bioconductor package edgeR (version 3.9) was used to analyze read count data and to identify differential expression levels (Benjamini–Hochberg method). We classified genes as being differentially expressed between genotypes when the differences in expression level between wildtype and homozygous mutant fish were significant at a false discovery rate (FDR) of 5%.

### Quantitative PCRs

One metal bead (Peqlab Biotechnologie) and 700 μl TRI Reagent (Sigma-Aldrich, St. Louis, USA) were added to the previously dissected and frozen eyes, forebrain, midbrain and hindbrain, followed by homogenization for 2 minutes at 30 Hz using the Qiagen tissue lyser. Total RNA was extracted from the different brain parts and eyes using the Direct-Zol RNA MiniPrep (Zymo Research, USA) according to the manufacturer’s protocol. 80ng-200ng of total RNA was transcribed into cDNA using the QuantiTect Reverse Transcription kit (Qiagen) with random hexamer primers. Each cDNA was further analyzed in duplicate in a 25ul volume using a SYBR green-containing mastermix in the Applied Biosystems (Thermo Fisher Scientific) mastercycler. Intron spanning qPCR primers were designed with the universal probe library software from Roche. Expression of each gene was normalized to *beta-actin*) transcript levels and fold changes were calculated.

**Table 1:**
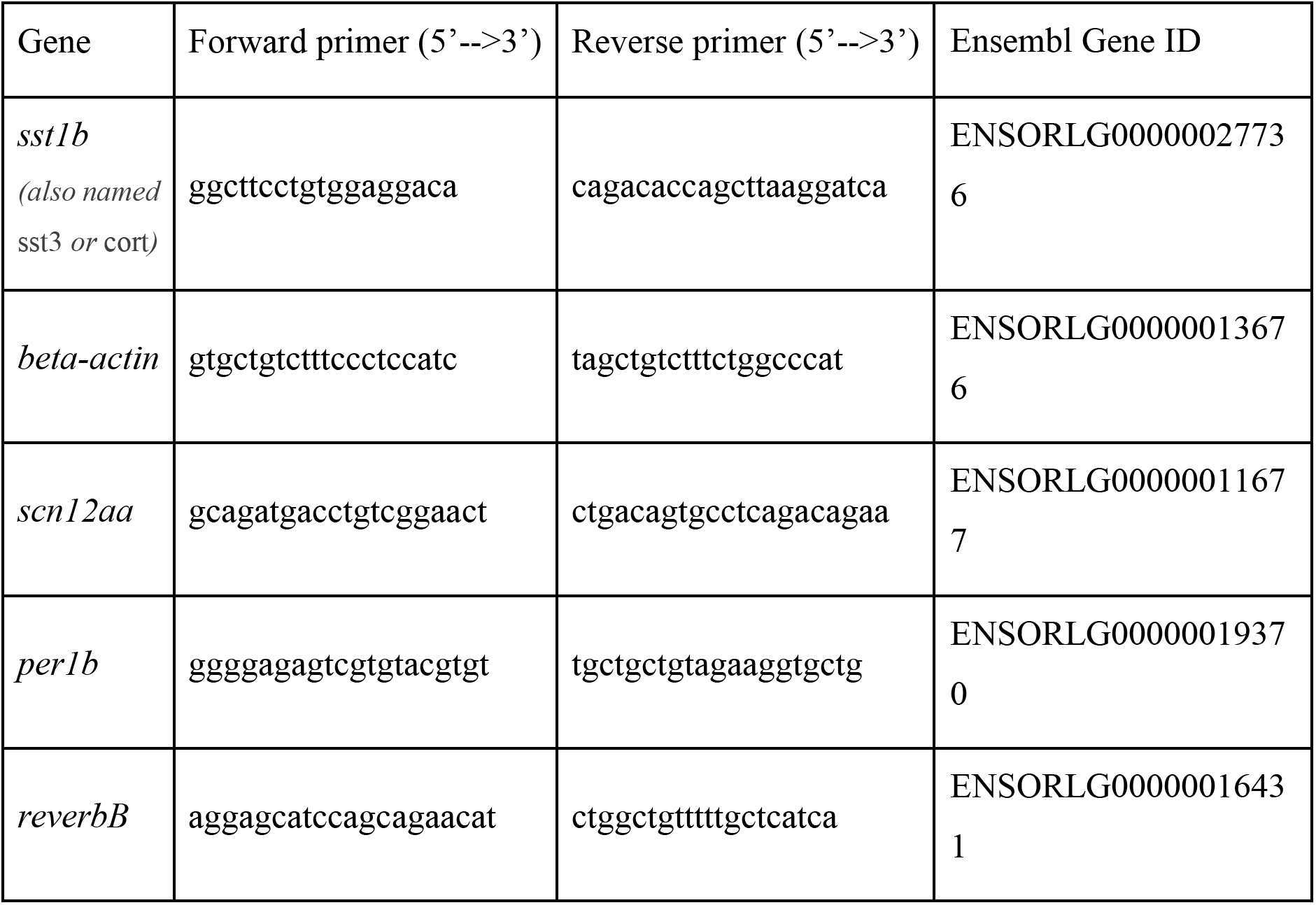
Forward and reverse primer sequences used for qPCR

### Cloning of medaka *gad2, scn12aa*

A BLAST search using the Zebrafish gad2 identified predicted homologous sequences, homology was confirmed by reciprocal back-blast (for *gad2*).

cDNAs were cloned following standard protocols, using Phusion Polymerase (NEB) and cloned in pJET1.2. Inserts were verified by sequencing.

Primers (5’-3’):

*gad2*

forward: atggcatctcacgggttctggtctc

reverse: ttatagatcatggccgaggcgttca

*scn12aa:*

forward :cgccatgcagagagaagattag

reverse: gacactcggtggtgaagattac

GenBank accession numbers: MT267535, MT239386, MT793601

### *In situ* hybridization (ISH) on adult brain sections

The generation of antisense digoxigenin (DIG)-labeled RNA probes and the ISH staining procedure on adult brain sections were performed as previously [14]. Briefly, adult fish (> 2 months-old) were anesthetized in fish water containing 0.2% tricaine, decapitated, and the brain dissected, fixed in ice-cold 4% PFA overnight, and subsequently stored in 100% methanol for at least 1 week prior to use. After rehydration in 1X PTW (PBS + 0.1% Tween-20), brains were digested with fresh 10 μg/ml Proteinase K (Merck, USA) for 35 minutes, followed by a 4 hours prehybridization step at 65°C with Hyb+ solution (50% Formamide, 5x SSC (pH=6.0), 0.1% Tween-20, 0.5 mg/ml torula (yeast) RNA, 50 μg/ml Heparin). Incubation with the specific DIG-RNA probe was made overnight at 65°C. Coronal whole-brain slices (100 μm thickness) were made using a vibrating blade microtome (Leica VT1000S, Leica Biosystems, Germany), blocked for > 1 hour in 10% sheep serum / 1x PTW, followed by incubation with 1:2000 dilution of anti-DIG-alkaline phosphatase-coupled antibody (Roche, Switzerland) in 10% sheep serum overnight. Detection of DIG-probes was made in staining buffer (in 10% polyvinyl alcohol) supplemented with nitro-blue tetrazolium (NBT) and 5-bromo-4-chloro-3’-indolyphosphate (BCIP) (Roche). In addition, the *gad2* probe was co-detected fluorescently as described in [74].

### Cortisol assay

#### Mechanical disturbance

Larvae were placed onto an orbital shaker (300rpm) for 5 minutes, subsequently sacrificed and snap-frozen in liquid nitrogen.

#### Cortisol extraction and measurement

The procedure for extracting cortisol from whole larvae was adapted from [75]. Briefly, the larvae were sacrificed and frozen in liquid nitrogen, and stored at −80°C until further processed. The samples were thawed on ice and 200ul sterile PBS was added. The samples were subsequently homogenized at RT for 1 minute at 30Hz (Tissue Lyser II, Qiagen). 20ul of each homogenate was removed for protein concentration measurements (Pierce BCA Protein Assay Kit, Thermo Scientific).

To the remaining 180ul, 1.4ml Ethyl acetate was added. The samples were subsequently centrifuged at 7000g for 15 minutes and the upper organic phase was collected within a glass vile. The samples were left to evaporate in a fume hood overnight. The dried samples were resuspended in 180ul ELISA Buffer (as provided by the Kit mentioned below).

The concentration of cortisol was measured using and enzyme-linked immunosorbent assay (ELISA) kit (500360, Cayman Chemical).

### Phylogenetic Analysis

Sequences were aligned using the MUSCLE alignment algorithm (https://www.ebi.ac.uk/Tools/msa/muscle/). The resulting alignment was used to generate a maximum likelihood (ML) tree by the IQ-TREE web server [76], visualized using iTol (https://itol.embl.de/) and exported as .eps file. Coloration and text formatting was performed in Adobe IllustratorCC2017. The accession numbers can be found in the respective metadata table.

### Statistics

The D’Agostino & Pearson normality test was used to asses data distribution. The F-test for comparison of variances for normally distributed data was used to assess the variance of the standard deviation. When comparing two normally distributed groups of data with the same standard deviation the unpaired t test was used. For comparing two groups of normally distributed data from with different standard deviations, the unpaired t test with Welch correction was used. The Mann-Whitney (M-W) test was used when comparing two non-normally distributed data sets. The software used was Prism version 8.

Data is presented as mean ± s.e.m. For all statistical tests used, a two-tailed *P* value was chosen: * *P* ≤ 0.05; ** *P* ≤ 0.01; *** *P* ≤ 0.001; **** *P* ≤ 0.0001

## Acknowledgements

We thank Andrij Belokurov and Margaryta Borisova for fish care, Andrew Straw and the members of the Tessmar-Raible, Raible, von Haeseler and Baier labs for helpful discussions, Mario Wullimann for advice on medaka neuroanatomy, Mirta Resetar for preparation of fish brains and RNA for DEseq analyses and Sven Schenk for sharing the mRNA sequencing protocol. Marin Čagelj contributed with the illustrations on Fig 1C, 4A, 5A.

## Supporting Information

**S1 Fig: relates to Fig 1,4.**
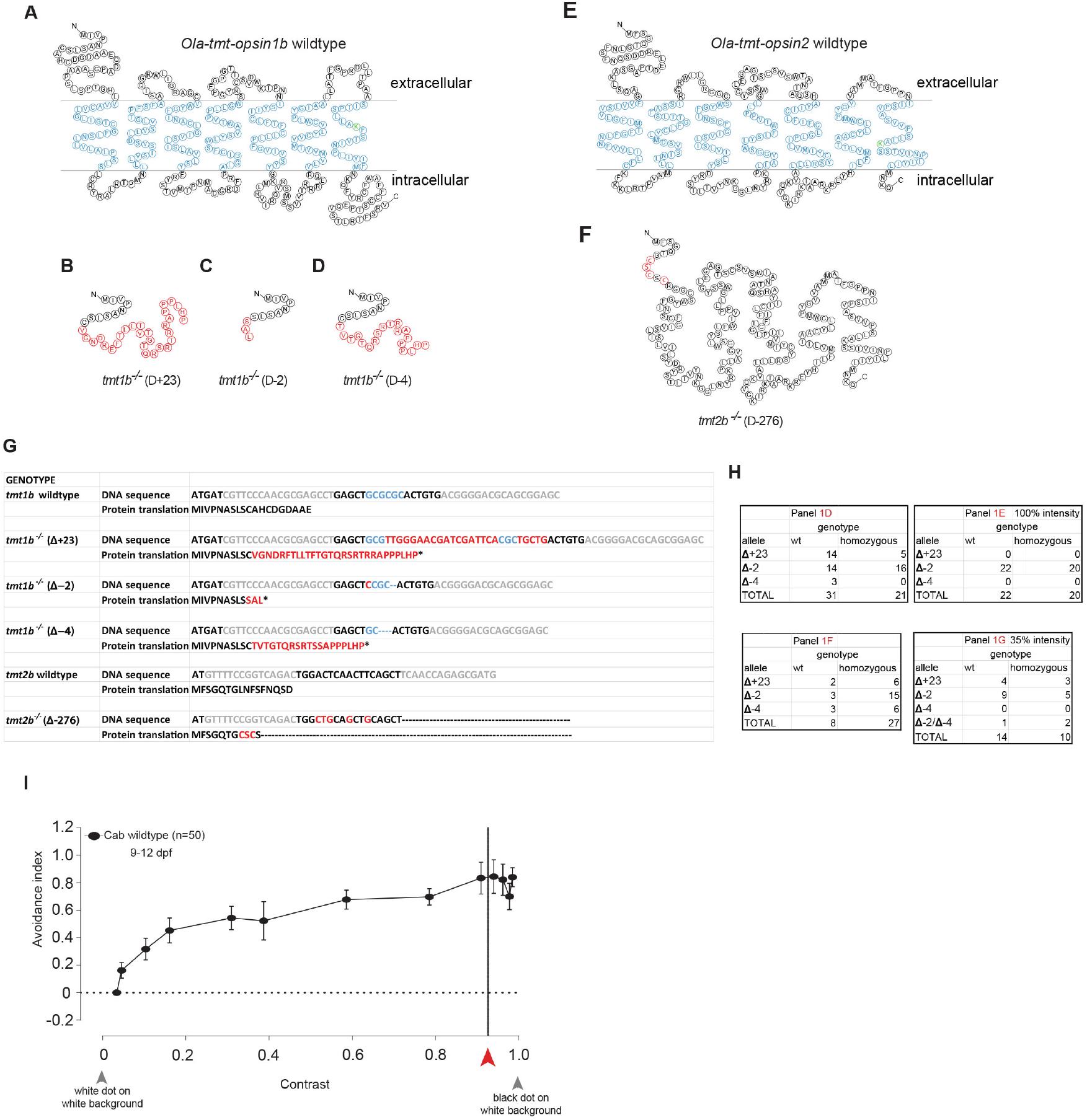
(**A-F**) Schematic of the predicted amino acid (aa) sequence for wildtype TMT-Opsin1b (A), wildtype TMT-Opsin2 (E), and corresponding recovered single TMT-Opsin1b (B-D) and TMT-Opsin2 (F) mutants. Blue aa: transmembrane domains; Green aa: Lys296 forming the chromophore’s Schiff base; Red aa: substituted amino acids. (**G**) Partial *Ola-tmt-opsin1b* and *Ola-tmt-opsin2* genomic DNA sequence and corresponding aa codon translation for wildtype and corresponding mutants that were used interchangeably along the study (grey: TALEN binding sites sequences; blue: recognition enzymatic site for BssHII; red: inserted nucleotides and substituted aa; *: stop codon). All the *tmt-opsin1b* mutants had in common the disruption of the BssHII recognition site (used for mutagenesis PCR confirmation) and a premature stop codon which led to a possible N-terminal truncated and hence null-functional protein. (**H**) Quantification of the number and identity of the wildtype and mutant allele’s siblings used in each trial. (**I**) Avoidance index levels of Cab wildtype larvae different contrast luminance levels between the moving dot (constant 20° size) and the white computer screen background. Red arrowhead indicated the contrast level measured at 35% light intensity. Note that at this contrast value, the level of avoidance is comparable to the avoidance levels revealed at higher contrasts. (Metadata: S7 Data)

**S2 Fig: relates to Fig 1.**
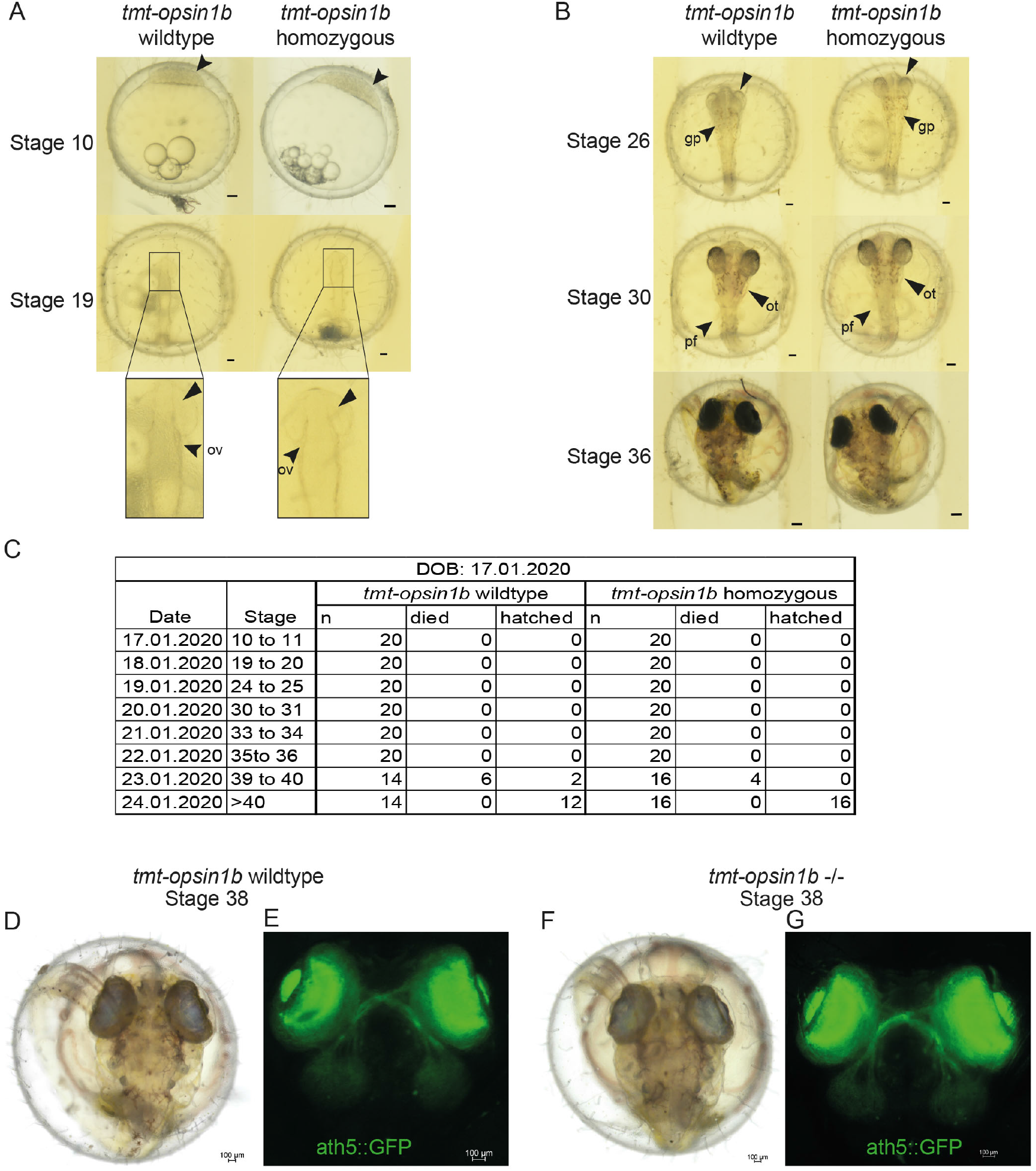
(**A,B**) Embryonic staging and development using morphological criteria for wildtype and *tmt1b* −/− mutant according to [33]. Some representative details: Stage 10 shows for both wildtype and mutant a thick blastoderm, with smaller inner cells. At Stage 19 both wildtype and mutant show the appearance of otic vesicles (ov), as indicated. Furthermore, a groove in the optic lobes can be observed. With Stage 26, both wildtype and mutant show that the choriodea start to cluster, as indicated with the arrow, and the guanophores (gp) are clearly visible. At Stage 30, the pectoral fin (pf) is apparent for both, as well as the otholits (ot). For Stage 36, the tip of the tail reaches the otic vesicle and the guanophores are now distributed from head to tail. Scale bar: 500μm (ov: otic vesicle, gp: guanophores, pf: pectoral fin, ot: otholits). (**C**) The table shows the total amount of embryos analyzed, with representatives depicted in A. Both wildtype and mutant each have an n=20. (**D-G**) Representative images of wildtype and *tmt-opsin1b* −/− mutants shortly before hatching, expressing GFP in the fish’s retinal ganglion cells shows no noticeable difference in axonal projections. Scale bars: 100μm

**S3 Fig: relates to Fig 1-4,7.**
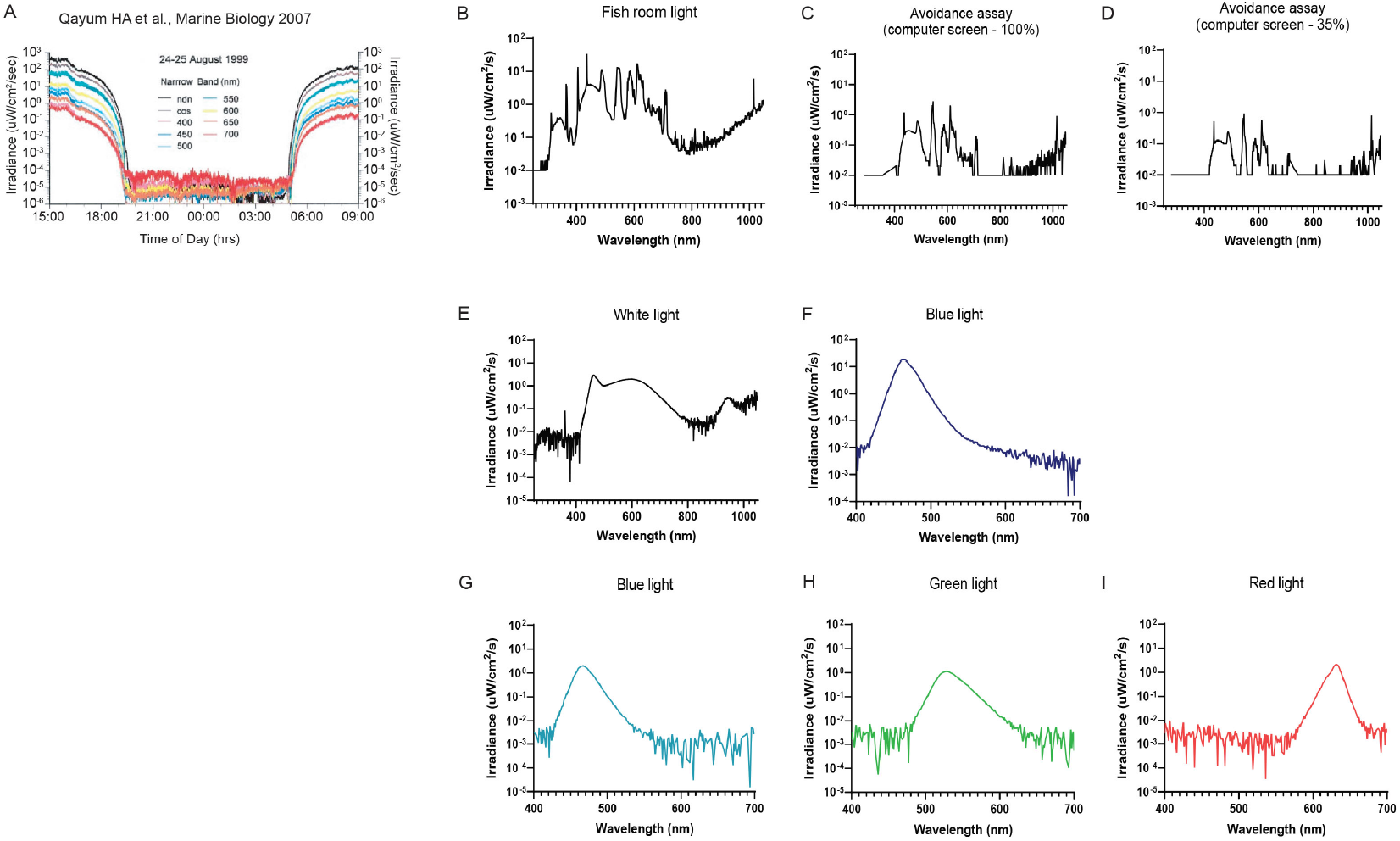
The units from our data are converted and plotted on logarithmic scale to facilitate comparisons with the published dataset. (**A**) Underwater light measurements for individual wavelengths between August 24-25, 1999 [77]. For calculations considering water turbidity also see [78]. (**B**) Light spectra of the fish room lighting, in which the fish were raised if not stated otherwise. (**C,D**) Light spectra of Fig 1I. (**E,F**) Light spectra of S4A Fig. (**G,H,I**) Light spectra of Fig 2I,K and M plotted as Irradiance.

**S4 Fig: relates to Fig 2.**
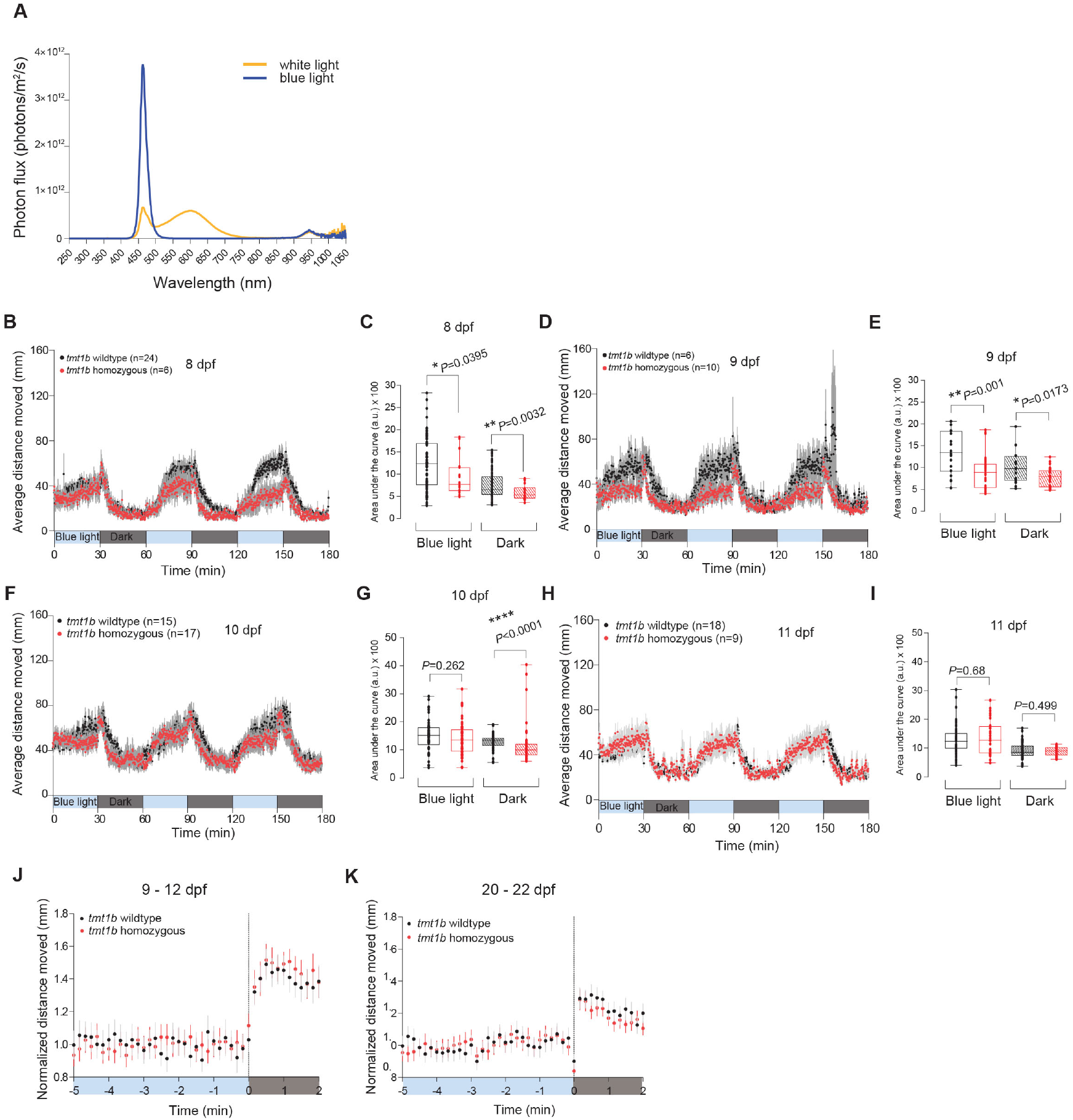
(**A**) Light spectra of the different LED lights used in the photokinesis assays (white light: yellow trace; Blue light: blue trace). (**B-I**) Average distance moved profile during alternating 30 minutes light / dark intervals and corresponding locomotor activity (as measured by the area under the curve) for 8 dpf (B,C), 9 dpf (D,E), 10 dpf (F,G) and 11 dpf (H,I) larvae‥ Each data point represents the mean (± s.e.m.) distance moved for the preceding 10 seconds. (**J,K**) Dark photokinesis response normalized to the average of the 5 minutes preceding darkness for larvae (J) and juvenile fish (K). All panels: (black*) tmt-opsin1b* wildtype and (red) *tmt-opsin1b* mutant. Colored boxes along the x-axis represent light condition. Box plots: single bar: median, box: first and third quartile, whiskers: min and max. a.u.: arbitrary units. (Metadata: S8 Data)

**S5 Fig: relates to Fig 3 and Fig 4.**
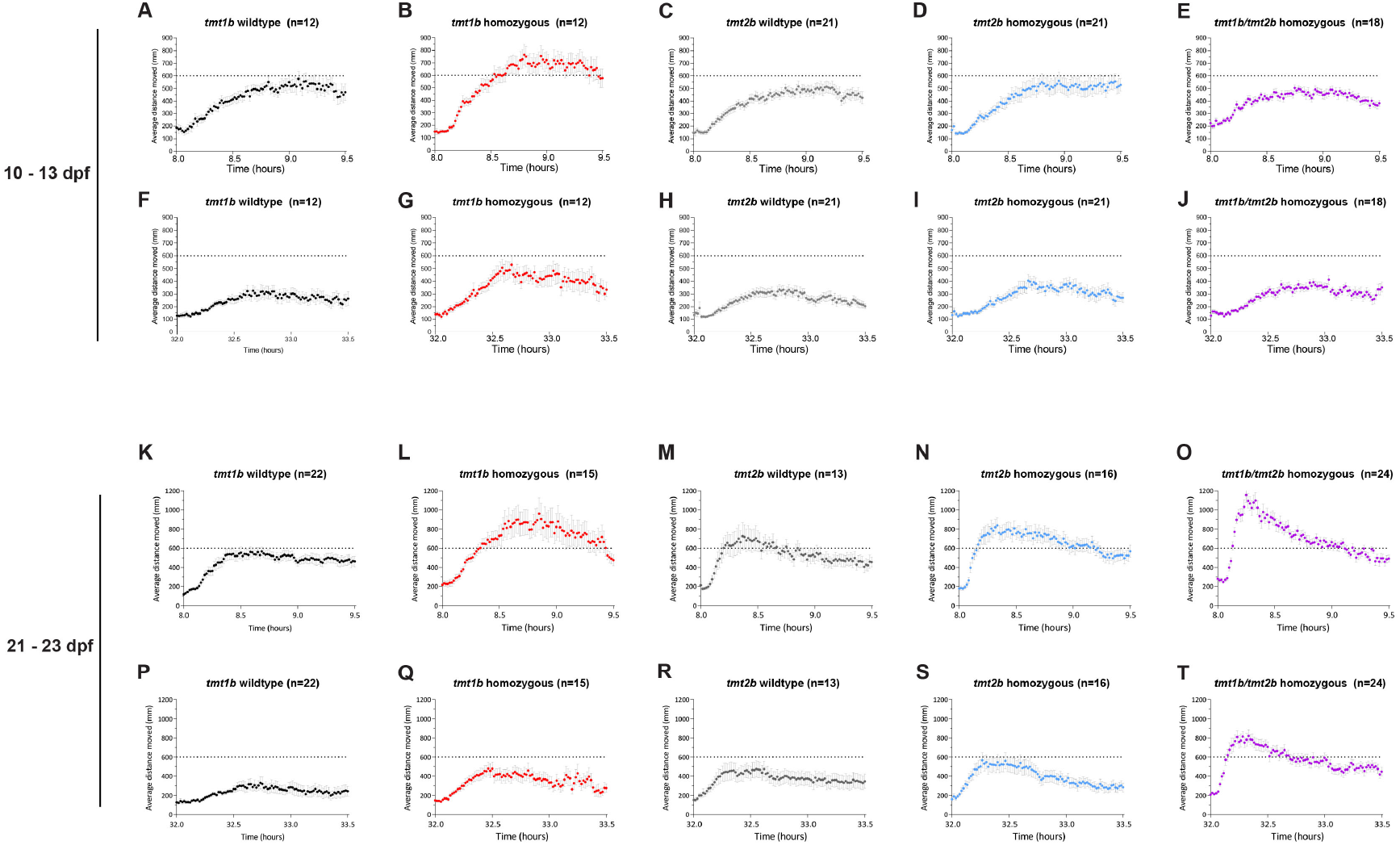
(**A-T**) Individual and higher time-resolved plots of average distance moved daytime shown for the first 1,5 hours of the first (A-E and K-O) and second day (F-J and P-T) of the experiment in larvae (A-J) and juvenile fish (K-T). Each data point represents the mean (± s.e.m.) distance moved for the preceding 1 minute. Metadata for A,B,F,G,K,L,P and Q : S3 Data Metadata for C,D,E,H,I,J,M,N,O,R,S and T : S4 Data

**S6 Fig: relates to Fig 6.**
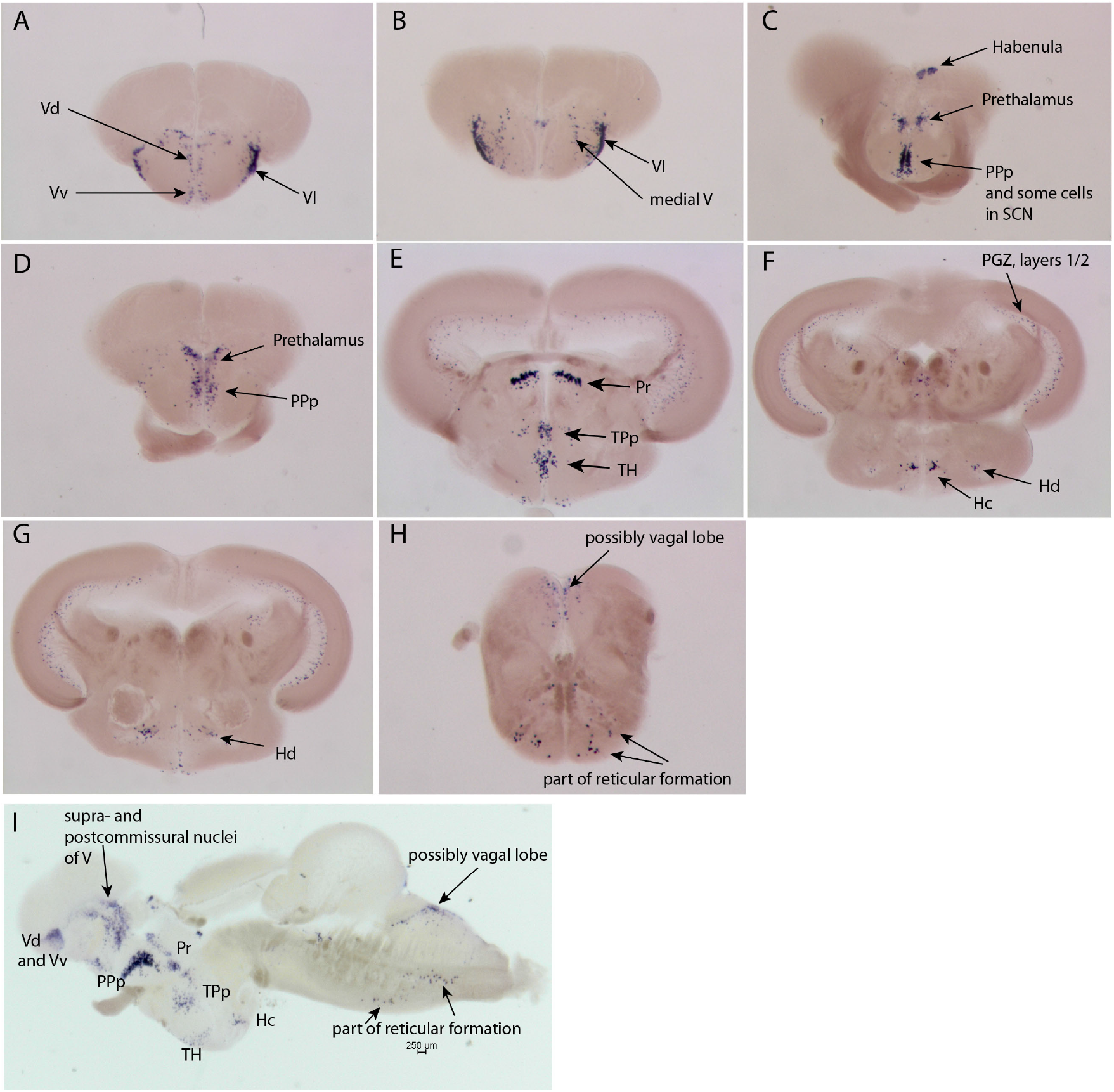
Anatomical annotation of *sst1b+* cells as revealed by *in situ* hybridisation in the adult medakafish brain, coronal/cross (**A-H**) and sagittal (**I**) sections. Abbreviations: (Hc) Caudal hypothalamus; (Hd) Dorsal hypothalamus; (PGZ) Periventricular gray zone; (Pr) Pretectum; (PPp) Posterior parvocellular preoptic nucleus; (SCN) Suprachiasmatic nucleus; (TH) Tuberal hypothalamus; (TPp) Periventricular posterior tuberculum; (Vd) Dorsal, (Vv) ventral, (Vl) lateral nucleus of (V) ventral telencephalic area.

**S7 Fig: relates to Fig 6.**
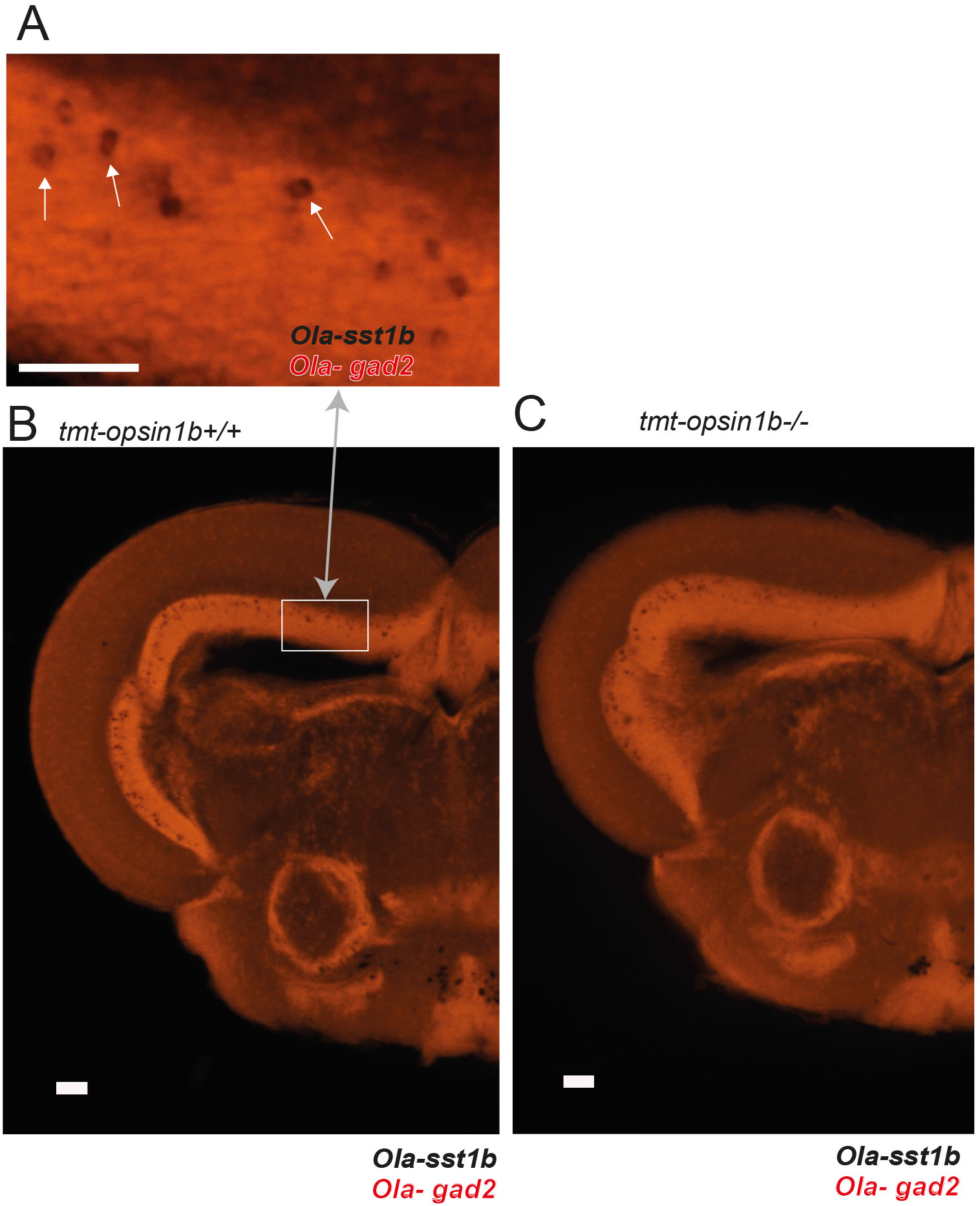
**(A)** *Sst1* co-expression with *gad2* in tectal neurons (arrows). **(B,C)** Representative images of *in situ* hybridizations for *sst1b* and *gad2* on coronal sections of the midbrain from *tmt-opsin1b* wildtype (B) and *tmt-opsin1b* homozygous (C) fish document no difference between wt and mutant fish. Red channel: *gad2*. Scale bars: 200μm

**S8 Fig: relates to Fig 6.**
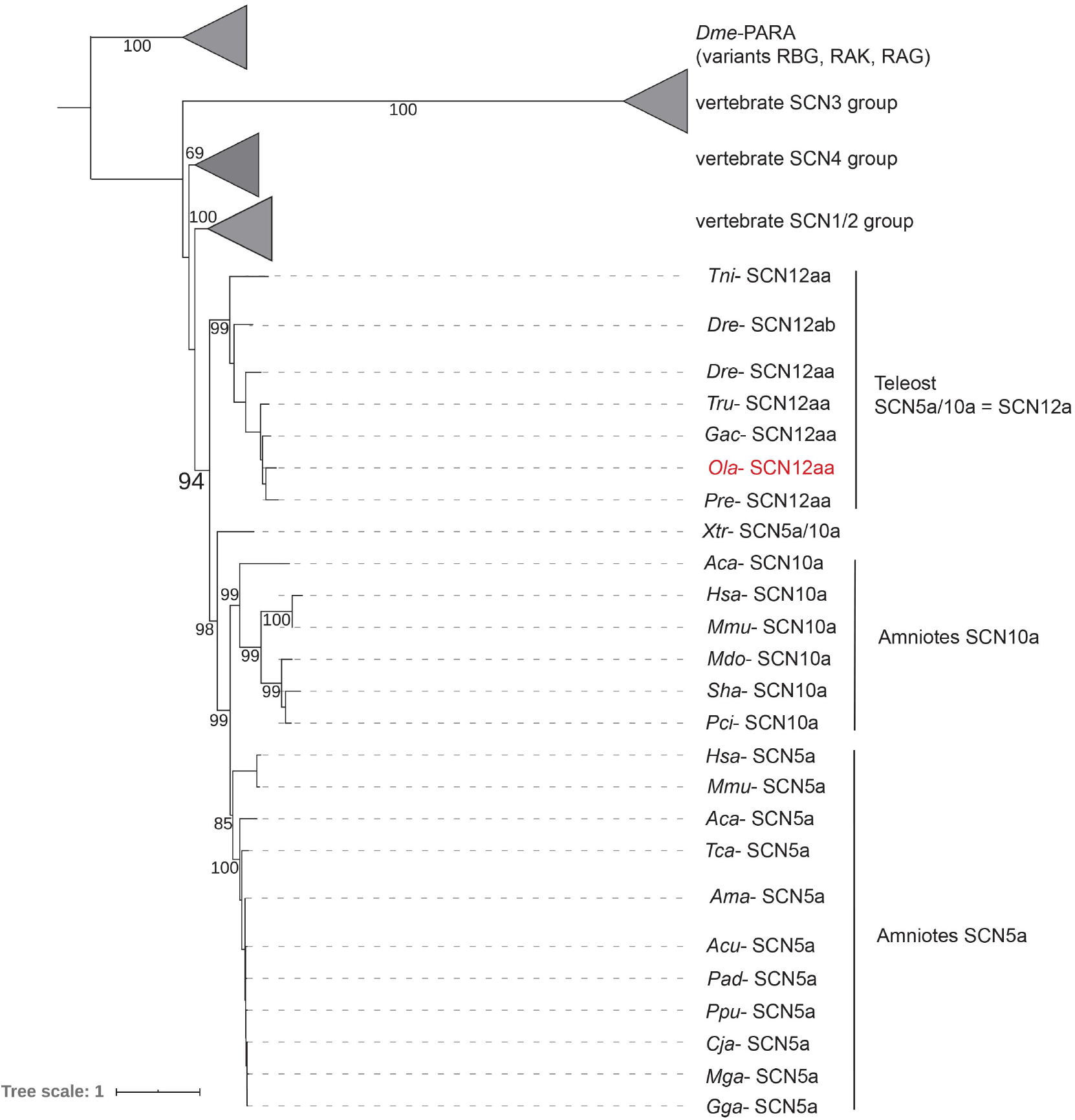
Maximum likelihood (ML) tree to resolve the orthology relationships of different voltage-gated sodium channel subunits across vertebrates. SCN12aa from Medaka fish is part of an ancestral fish/amphibian group, which duplicated in amniotes giving rise to two distinct groups, SCN5a and SCN10a. Variants of the *Drosophila* voltage-gated sodium channel PARA are used as outgroup. For accession numbers of the proteins used for alignment and tree construction: S9 Data.

**S9 Fig: relates to Fig 6.**
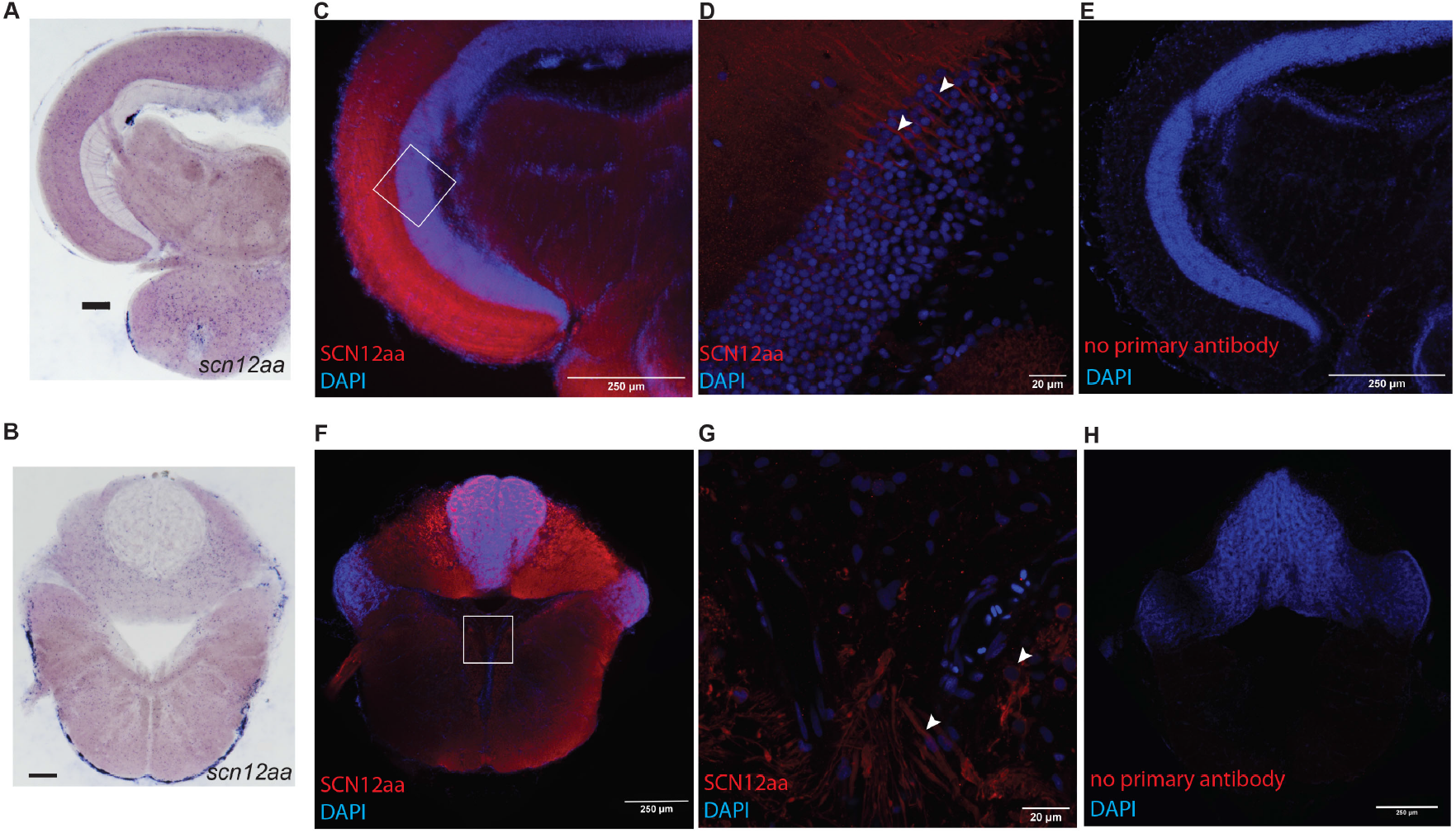
(**A,B**) In situ hybridization for *scn12aa* performed on coronal sections of midbrain (A) and hindbrain (B) from Cab wildtype fish. Scale bar: 50 μm. (**C-G**) Confocal images of brains stained for SCN12aa, using an antibody raised against *Hsa*-Scn5a (red) and DAPI as nuclear counterstain (blue) on coronal sections of midbrain (C,D) and hindbrain (F,G) from Cab wildtype fish. White box: Region corresponding to the 40x magnification. White arrowheads: Representative positively stained neuronal projections and cell bodies. **(E,H)** Control slices with equal treatment as for C,F, but no primary antibody added.

**S1 Video: relates to Fig 1E**

Example of a 10 dpf *tmt-opsin1b* wildtype larvae performing the avoidance assay during the ascending size dot assay. Large dots elicit a distinct avoidance behavior.

**S2 Video: relates to Fig 1E**

Example of a 10 dpf *tmt-opsin1b* single homozygous larvae performing the avoidance assay during the ascending size dot assay. Large dots elicit a distinct avoidance behavior. Note that the displayed number of avoidances is higher when compared with wildtype larvae.

## References

1. Posch T, Freyhoff A, Uhlmann T. Das Ende der Nacht: Die globale Lichtverschmutzung und ihre Folgen. 1st edition ed. Weinheim: Wiley VCH; 2010. 154 p.

2. Hunt DM, Hankins MW, Collin SP, Marshall NJ. Evolution of visual and non-visual pigments. New York: Springer; 2014. 276 p.

3. Bedrosian TA, Nelson RJ. Timing of light exposure affects mood and brain circuits. Transl Psychiatry. 2017;7(1):e1017. Epub 2017/02/01. doi: 10.1038/tp.2016.262. PubMed PMID: 28140399; PubMed Central PMCID: PMCPMC5299389.

4. Wahl S, Engelhardt M, Schaupp P, Lappe C, Ivanov IV. The inner clock-Blue light sets the human rhythm. J Biophotonics. 2019;12(12):e201900102. Epub 2019/08/23. doi: 10.1002/jbio.201900102. PubMed PMID: 31433569; PubMed Central PMCID: PMCPMC7065627.

5. Fernandez DC, Fogerson PM, Lazzerini Ospri L, Thomsen MB, Layne RM, Severin D, et al. Light Affects Mood and Learning through Distinct Retina-Brain Pathways. Cell. 2018;175(1):71–84 e18. Epub 2018/09/04. doi: 10.1016/j.cell.2018.08.004. PubMed PMID: 30173913; PubMed Central PMCID: PMCPMC6190605.

6. Zhu H, Wang N, Yao L, Chen Q, Zhang R, Qian J, et al. Moderate UV Exposure Enhances Learning and Memory by Promoting a Novel Glutamate Biosynthetic Pathway in the Brain. Cell. 2018;173(7):1716–27 e17. Epub 2018/05/22. doi: 10.1016/j.cell.2018.04.014. PubMed PMID: 29779945.

7. Dulcis D, Jamshidi P, Leutgeb S, Spitzer NC. Neurotransmitter switching in the adult brain regulates behavior. Science. 2013;340(6131):449–53. Epub 2013/04/27. doi: 10.1126/science.1234152. PubMed PMID: 23620046.

8. Dawson A, King VM, Bentley GE, Ball GF. Photoperiodic control of seasonality in birds. J Biol Rhythms. 2001;16(4):365–80. Epub 2001/08/17. doi: 10.1177/074873001129002079. PubMed PMID: 11506381.

9. Currie SP, Doherty GH, Sillar KT. Deep-brain photoreception links luminance detection to motor output in Xenopus frog tadpoles. Proc Natl Acad Sci U S A. 2016;113(21):6053–8. Epub 2016/05/12. doi: 10.1073/pnas.1515516113. PubMed PMID: 27166423; PubMed Central PMCID: PMCPMC4889350.

10. von Frisch K. Beiträge zur Physiologie der Pigmentzellen in der Fischhaut. Pflüg Arch ges Physiol. 1911;138:319–87.

11. Foster RG, Follett BK, Lythgoe JN. Rhodopsin-like sensitivity of extra-retinal photoreceptors mediating the photoperiodic response in quail. Nature. 1985;313(5997):50–2. Epub 1985/01/03. PubMed PMID: 3965970.

12. Halford S, Pires SS, Turton M, Zheng L, Gonzalez-Menendez I, Davies WL, et al. VA opsin-based photoreceptors in the hypothalamus of birds. Curr Biol. 2009;19(16):1396–402. Epub 2009/08/12. doi: 10.1016/j.cub.2009.06.066. PubMed PMID: 19664923.

13. Fernandes AM, Fero K, Arrenberg AB, Bergeron SA, Driever W, Burgess HA. Deep brain photoreceptors control light-seeking behavior in zebrafish larvae. Curr Biol. 2012;22(21):2042–7. Epub 2012/09/25. doi: 10.1016/j.cub.2012.08.016. PubMed PMID: 23000151; PubMed Central PMCID: PMC3494761.

14. Fischer RM, Fontinha BM, Kirchmaier S, Steger J, Bloch S, Inoue D, et al. Co-expression of VAL- and TMT-opsins uncovers ancient photosensory interneurons and motorneurons in the vertebrate brain. PLoS Biol. 2013;11(6):e1001585. doi: 10.1371/journal.pbio.1001585.

15. Davies WIL, Tamai TK, Zhen L, Fu JK, Rihel J, Foster RG, et al. An extended family of novel vertebrate photopigments is widely expressed and displays a diversity of function. Genome Res. 2015;25(11):1666–79. doi: 10.1101/gr.189886.115. PubMed PMID: WOS:000364355600008.

16. Jenkins A, Munoz M, Tarttelin EE, Bellingham J, Foster RG, Hankins MW. VA opsin, melanopsin, and an inherent light response within retinal interneurons. Current Biology : CB. 2003;13(15):1269–78. Epub 2003/08/09. PubMed PMID: 12906786.

17. Davies WI, Zheng L, Hughes S, Tamai TK, Turton M, Halford S, et al. Functional diversity of melanopsins and their global expression in the teleost retina. Cell Mol Life Sci. 2011;68(24):4115–32. Epub 2011/08/13. doi: 10.1007/s00018-011-0785-4. PubMed PMID: 21833582.

18. Nakane Y, Shimmura T, Abe H, Yoshimura T. Intrinsic photosensitivity of a deep brain photoreceptor. Curr Biol. 2014;24(13):R596–7. Epub 2014/07/09. doi: 10.1016/j.cub.2014.05.038. PubMed PMID: 25004360.

19. Brunt EEV, Shepherd MD, Wall JR, Ganong WF, Clegg MT. Penetration of light into the brain of mammals. Ann N Y Acad Sci. 1964;117(1):217–24. doi: 10.1111/j.1749-6632.1964.tb48177.x.

20. Kokel D, Dunn TW, Ahrens MB, Alshut R, Cheung CY, Saint-Amant L, et al. Identification of nonvisual photomotor response cells in the vertebrate hindbrain. The Journal of Neuroscience. 2013;33(9):3834–43. Epub 2013/03/01. doi: 10.1523/JNEUROSCI.3689-12.2013. PubMed PMID: 23447595; PubMed Central PMCID: PMC3600642.

21. Friedmann D, Hoagland A, Berlin S, Isacoff EY. A spinal opsin controls early neural activity and drives a behavioral light response. Curr Biol. 2015;25(1):69–74. Epub 2014/12/09. doi: 10.1016/j.cub.2014.10.055. PubMed PMID: 25484291; PubMed Central PMCID: PMC4286461.

22. Horstick EJ, Bayleyen Y, Sinclair JL, Burgess HA. Search strategy is regulated by somatostatin signaling and deep brain photoreceptors in zebrafish. BMC Biol. 2017;15. doi: ARTN 4 10.1186/s12915-016-0346-2. PubMed PMID: WOS:000392962100002.

23. Matos-Cruz V, Blasic J, Nickle B, Robinson PR, Hattar S, Halpern ME. Unexpected diversity and photoperiod dependence of the zebrafish melanopsin system. PLoS ONE. 2011;6(9):e25111. Epub 2011/10/04. doi: 10.1371/journal.pone.0025111. PubMed PMID: 21966429; PubMed Central PMCID: PMCPMC3178608.

24. Cavallari N, Frigato E, Vallone D, Frohlich N, Lopez-Olmeda JF, Foa A, et al. A blind circadian clock in cavefish reveals that opsins mediate peripheral clock photoreception. PLoS Biol. 2011;9(9):e1001142. Epub 2011/09/13. doi: 10.1371/journal.pbio.1001142. PubMed PMID: 21909239; PubMed Central PMCID: PMC3167789.

25. Moore HA, Whitmore D. Circadian Rhythmicity and Light Sensitivity of the Zebrafish Brain. PLoS ONE. 2014;9(1). doi: ARTN e86176 10.1371/journal.pone.0086176. PubMed PMID: WOS:000330283100121.

26. Nakane Y, Ikegami K, Iigo M, Ono H, Takeda K, Takahashi D, et al. The saccus vasculosus of fish is a sensor of seasonal changes in day length. Nat Commun. 2013;4. doi: ARTN 2108 10.1038/ncomms3108. PubMed PMID: WOS:000323715700003.

27. Ramirez MD, Pairett AN, Pankey MS, Serb JM, Speiser DI, Swafford AJ, et al. The Last Common Ancestor of Most Bilaterian Animals Possessed at Least Nine Opsins. Genome Biol Evol. 2016;8(12):3640–52. doi: 10.1093/gbe/evw248. PubMed PMID: 28172965; PubMed Central PMCID: PMCPMC5521729.

28. Blackshaw S, Snyder SH. Encephalopsin: a novel mammalian extraretinal opsin discretely localized in the brain. J Neurosci. 1999;19(10):3681–90.

29. Nissila J, Manttari S, Sarkioja T, Tuominen H, Takala T, Timonen M, et al. Encephalopsin (OPN3) protein abundance in the adult mouse brain. J Comp Physiol A Neuroethol Sens Neural Behav Physiol. 2012;198(11):833–9. Epub 2012/09/20. doi: 10.1007/s00359-012-0754-x. PubMed PMID: 22991144; PubMed Central PMCID: PMC3478508.

30. Awaji M, Hanyu I. Seasonal Changes in Ovarian Response to Photoperiods in Orange-Red Type Medaka : Reproductive Biology. Zoological Science. 1989;6:943–50.

31. Koger CS, Teh SJ, Hinton DE. Variations of light and temperature regimes and resulting effects on reproductive parameters in medaka (Oryzias latipes). Biol Reprod. 1999;61(5):1287–93. Epub 1999/10/21. PubMed PMID: 10529276.

32. Sakai K, Yamashita T, Imamoto Y, Shichida Y. Diversity of Active States in TMT Opsins. PLoS ONE. 2015;10(10):e0141238. doi: 10.1371/journal.pone.0141238. PubMed PMID: 26491964; PubMed Central PMCID: PMCPMC4619619.

33. Kinoshita M, Murata K, Naruse K, Tanaka M. Medaka: Biology, Management, and Experimental Protocols: Wiley-Blackwell; 2009.

34. Souren M, Martinez-Morales JR, Makri P, Wittbrodt B, Wittbrodt J. A global survey identifies novel upstream components of the Ath5 neurogenic network. Genome Biol. 2009;10(9):R92. Epub 2009/09/09. doi: 10.1186/gb-2009-10-9-r92. PubMed PMID: 19735568; PubMed Central PMCID: PMCPMC2768981.

35. Del Bene F, Ettwiller L, Skowronska-Krawczyk D, Baier H, Matter JM, Birney E, et al. In vivo validation of a computationally predicted conserved Ath5 target gene set. PLoS Genet. 2007;3(9):1661–71. Epub 2007/09/26. doi: 10.1371/journal.pgen.0030159. PubMed PMID: 17892326; PubMed Central PMCID: PMC1988851.

36. Ewert JP. Neural Mechanisms of Prey-Catching and Avoidance Behavior in Toad (Bufo-Bufo L). Brain Behavior and Evolution. 1970;3(1-4):36-&. doi: Doi 10.1159/000125462. PubMed PMID: WOS:A1970I326400005.

37. Dong W, Lee RH, Xu H, Yang S, Pratt KG, Cao V, et al. Visual Avoidance in *Xenopus* Tadpoles Is Correlated With the Maturation of Visual Responses in the Optic Tectum. J Neurophysiol. 2009;101(2):803–15. doi: 10.1152/jn.90848.2008. PubMed PMID: WOS:000263120300027.

38. Khakhalin AS, Koren D, Gu J, Xu H, Aizenman CD. Excitation and inhibition in recurrent networks mediate collision avoidance in *Xenopus* tadpoles. Eur J Neurosci. 2014;40(6):2948–62. doi: 10.1111/ejn.12664. PubMed PMID: 24995793.

39. Semmelhack JL, Donovan JC, Thiele TR, Kuehn E, Laurell E, Baier H. A dedicated visual pathway for prey detection in larval zebrafish. Elife. 2014;3. doi: ARTN e04878 10.7554/eLife.04878. PubMed PMID: WOS:000346170300001.

40. Barker AJ, Baier H. Sensorimotor Decision Making in the Zebrafish Tectum. Curr Biol. 2015;25(21):2804–14. doi: 10.1016/j.cub.2015.09.055. PubMed PMID: WOS:000364262500022.

41. Yilmaz M, Meister M. Rapid Innate Defensive Responses of Mice to Looming Visual Stimuli. Curr Biol. 2013;23(20):2011–5. doi: 10.1016/j.cub.2013.08.015. PubMed PMID: WOS:000326317300022.

42. Preuss T, Osei-Bonsu PE, Weiss SA, Wang C, Faber DS. Neural representation of object approach in a decision-making motor circuit. J Neurosci. 2006;26(13):3454–64. doi: 10.1523/Jneurosci.5259-05.2006. PubMed PMID: WOS:000236363400010.

43. Fraenkel GS, Gunn DL. The Orientation of Animals, Kineses, Taxes and Compass Reactions‥ New York: Dover Publications Inc.; 1961.

44. Van Veen T, Hartwig H, Müller K. Light-dependent motor activity and photonegative behavior in the eel (Anguilla anguilla L.). Journal of comparative physiology. 1976;111(2):209–19.

45. Young J. The photoreceptors of lampreys: I. Light-sensitive fibres in the lateral line nerves. J Exp Biol. 1935;12(3):229–38.

46. Burgess HA, Granato M. Modulation of locomotor activity in larval zebrafish during light adaptation. J Exp Biol. 2007;210(Pt 14):2526–39. doi: 10.1242/jeb.003939. PubMed PMID: 17601957.

47. Prober DA, Rihel J, Onah AA, Sung RJ, Schier AF. Hypocretin/orexin overexpression induces an insomnia-like phenotype in zebrafish. J Neurosci. 2006;26(51):13400–10. doi: 10.1523/JNEUROSCI.4332-06.2006. PubMed PMID: 17182791.

48. Sugihara T, Nagata T, Mason B, Koyanagi M, Terakita A. Absorption Characteristics of Vertebrate Non-Visual Opsin, Opn3. PLoS ONE. 2016;11(8):e0161215. doi: 10.1371/journal.pone.0161215. PubMed PMID: 27532629; PubMed Central PMCID: PMCPMC4988782.

49. Kato M, Sugiyama T, Sakai K, Yamashita T, Fujita H, Sato K, et al. Two Opsin 3-Related Proteins in the Chicken Retina and Brain: A TMT-Type Opsin 3 Is a Blue-Light Sensor in Retinal Horizontal Cells, Hypothalamus, and Cerebellum. PLoS ONE. 2016;11(11):e0163925. doi: 10.1371/journal.pone.0163925. PubMed PMID: 27861495; PubMed Central PMCID: PMCPMC5115664.

50. Naumann EA, Fitzgerald JE, Dunn TW, Rihel J, Sompolinsky H, Engert F. From Whole-Brain Data to Functional Circuit Models: The Zebrafish Optomotor Response. Cell. 2016;167(4):947–60 e20. Epub 2016/11/05. doi: 10.1016/j.cell.2016.10.019. PubMed PMID: 27814522; PubMed Central PMCID: PMCPMC5111816.

51. Kim DH, Kim J, Marques JC, Grama A, Hildebrand DGC, Gu W, et al. Pan-neuronal calcium imaging with cellular resolution in freely swimming zebrafish. Nat Methods. 2017;14(11):1107–14. Epub 2017/09/12. doi: 10.1038/nmeth.4429. PubMed PMID: 28892088.

52. Fero K, Yokogawa T, Burgess HA. The Behavioral Repertoire of Larval Zebrafish. In: Kalueff AV, Cachat JM, editors. Zebrafish Models in Neurobehavioral Research. Totowa, NJ: Humana Press; 2011. p. 249–91.

53. Carvalho PSM, Noltie DB, Tillitt DE. Ontogenetic improvement of visual function in the medaka Oryzias latipes based on an optomotor testing system for larval and adult fish. Anim Behav. 2002;64:1–10. doi: 10.1006/anbe.2002.3028. PubMed PMID: WOS:000178657200001.

54. Cuesta IH, Lahiri K, Lopez-Olmeda JF, Loosli F, Foulkes NS, Vallone D. Differential maturation of rhythmic clock gene expression during early development in medaka (Oryzias latipes). Chronobiol Int. 2014;31(4):468–78. Epub 2014/01/25. doi: 10.3109/07420528.2013.856316. PubMed PMID: 24456338.

55. Robinson MD, McCarthy DJ, Smyth GK. edgeR: a Bioconductor package for differential expression analysis of digital gene expression data. Bioinformatics. 2010;26(1):139–40. Epub 2009/11/17. doi: 10.1093/bioinformatics/btp616. PubMed PMID: 19910308; PubMed Central PMCID: PMCPMC2796818.

56. Raj B, Wagner DE, McKenna A, Pandey S, Klein AM, Shendure J, et al. Simultaneous single-cell profiling of lineages and cell types in the vertebrate brain. Nat Biotechnol. 2018;36(5):442–50. Epub 2018/04/03. doi: 10.1038/nbt.4103. PubMed PMID: 29608178; PubMed Central PMCID: PMCPMC5938111.

57. Nakayama T, Shimmura T, Shinomiya A, Okimura K, Takehana Y, Furukawa Y, et al. Seasonal regulation of the lncRNA LDAIR modulates self-protective behaviours during the breeding season. Nat Ecol Evol. 2019;3(5):845–52. Epub 2019/04/10. doi: 10.1038/s41559-019-0866-6. PubMed PMID: 30962562.

58. Song Y-H, Hwang Y-S, Kim K, Lee H-R, Kim J-H, Maclachlan C, et al. Somatostatin enhances visual processing and perception by suppressing excitatory inputs to parvalbumin-positive interneurons in V1. Science advances. 2020;6(17):eaaz0517–eaaz. doi: 10.1126/sciadv.aaz0517. PubMed PMID: 32494634.

59. Isoe Y, Konagaya Y, Yokoi S, Kubo T, Takeuchi H. Ontogeny and Sexual Differences in Swimming Proximity to Conspecifics in Response to Visual Cues in Medaka Fish. Zoolog Sci. 2016;33(3):246–54. Epub 2016/06/09. doi: 10.2108/zs150213. PubMed PMID: 27268978.

60. Pinto JG, Hornby KR, Jones DG, Murphy KM. Developmental changes in GABAergic mechanisms in human visual cortex across the lifespan. Front Cell Neurosci. 2010;4:16. Epub 2010/07/02. doi: 10.3389/fncel.2010.00016. PubMed PMID: 20592950; PubMed Central PMCID: PMCPMC2893712.

61. Siu CR, Beshara SP, Jones DG, Murphy KM. Development of Glutamatergic Proteins in Human Visual Cortex across the Lifespan. J Neurosci. 2017;37(25):6031–42. Epub 2017/05/31. doi: 10.1523/JNEUROSCI.2304-16.2017. PubMed PMID: 28554889; PubMed Central PMCID: PMCPMC6596503.

62. Dahlgren CP, Eggleston DB. Ecological Processes Underlying Ontogenetic Habitat Shifts in a Coral Reef Fish. Ecology. 2000;81(8):2227–40. doi: 10.2307/177110.

63. Nunn AD, Tewson LH, Cowx IG. The foraging ecology of larval and juvenile fishes. Reviews in Fish Biology and Fisheries. 2012;22(2):377–408. doi: 10.1007/s11160-011-9240-8.

64. Bannister S, Antonova O, Polo A, Lohs C, Hallay N, Valinciute A, et al. TALE Nucleases mediate efficient, heritable genome modifications in the marine annelid *Platynereis dumerilii*. Genetics. 2014;197(1):19–31. doi: 10.1534/genetics.112.148254.

65. Pende M, Vadiwala K, Schmidbaur H, Stockinger AW, Murawala P, Saghafi S, et al. A versatile depigmentation, clearing, and labeling method for exploring nervous system diversity. Sci Adv. 2020;6(22):eaba0365. Epub 2020/06/12. doi: 10.1126/sciadv.aba0365. PubMed PMID: 32523996; PubMed Central PMCID: PMCPMC7259959.

66. Matsunaga W, Watanabe E. Habituation of medaka (Oryzias latipes) demonstrated by open-field testing. Behav Processes. 2010;85(2):142–50. doi: 10.1016/j.beproc.2010.06.019. PubMed PMID: WOS:000282121400008.

67. Michelson AA. Studies in optics: Chicago, Ill. : The University of Chicago Press; 1927.

68. Beck JC, Gilland E, Tank DW, Baker R. Quantifying the ontogeny of optokinetic and vestibuloocular behaviors in zebrafish, medaka, and goldfish. J Neurophysiol. 2004;92(6):3546–61. doi: 10.1152/jn.00311.2004. PubMed PMID: WOS:000225164800038.

69. Kokel D, Bryan J, Laggner C, White R, Cheung CYJ, Mateus R, et al. Rapid behavior-based identification of neuroactive small molecules in the zebrafish. Nat Chem Biol. 2010;6(3):231–7. doi: 10.1038/Nchembio.307. PubMed PMID: WOS:000274565100016.

70. Lopez-Olmeda JF, Tartaglione EV, de la Iglesia HO, Sanchez-Vazquez FJ. Feeding entrainment of food-anticipatory activity and per1 expression in the brain and liver of zebrafish under different lighting and feeding conditions. Chronobiol Int. 2010;27(7):1380–400. Epub 2010/08/28. doi: 10.3109/07420528.2010.501926. PubMed PMID: 20795882.

71. Anken R, Bourrat F. Brain atlas of the Medakafish *Oryzias latipes*. Paris: Institut national de la recherche agronomique; 1998.

72. Sedlazeck FJ, Rescheneder P, von Haeseler A. NextGenMap: fast and accurate read mapping in highly polymorphic genomes. Bioinformatics. 2013;29(21):2790–1. doi: 10.1093/bioinformatics/btt468. PubMed PMID: 23975764.

73. Liao Y, Smyth GK, Shi W. featureCounts: an efficient general purpose program for assigning sequence reads to genomic features. Bioinformatics. 2014;30(7):923–30. doi: 10.1093/bioinformatics/btt656. PubMed PMID: WOS:000334078300005.

74. Tessmar-Raible K, Steinmetz PR, Snyman H, Hassel M, Arendt D. Fluorescent two-color whole mount in situ hybridization in Platynereis dumerilii (Polychaeta, Annelida), an emerging marine molecular model for evolution and development. Biotechniques. 2005;39(4):460, 2, 4. Epub 2005/10/21. PubMed PMID: 16235555.

75. Bai Y, Liu H, Huang B, Wagle M, Guo S. Identification of environmental stressors and validation of light preference as a measure of anxiety in larval zebrafish. BMC Neurosci. 2016;17(1):63. doi: 10.1186/s12868-016-0298-z.

76. Trifinopoulos J, Nguyen LT, von Haeseler A, Minh BQ. W-IQ-TREE: a fast online phylogenetic tool for maximum likelihood analysis. Nucleic Acids Res. 2016;44(W1):W232–5. Epub 2016/04/17. doi: 10.1093/nar/gkw256. PubMed PMID: 27084950; PubMed Central PMCID: PMCPMC4987875.

77. Qayum HA, Klimley AP, Newton R, Richert JE. Broad-band versus narrow-band irradiance for estimating latitude by archival tags. Marine Biology. 2007;151(2):467–81. doi: 10.1007/s00227-006-0514-y.

78. Kiang NY, Siefert J, Govindjee, Blankenship RE. Spectral signatures of photosynthesis. I. Review of Earth organisms. Astrobiology. 2007;7(1):222–51. Epub 2007/04/05. doi: 10.1089/ast.2006.0105. PubMed PMID: 17407409.

